# Systematic review and meta-analysis of bulk RNAseq studies in human Alzheimer’s disease brain tissue

**DOI:** 10.1101/2024.11.07.622520

**Authors:** Bernardo Aguzzoli Heberle, Kristin L. Fox, Lucas Lobraico Libermann, Sophia Ronchetti Martins Xavier, Guilherme Tarnowski Dallarosa, Rhaná Carolina Santos, David W. Fardo, Thiago Wendt Viola, Mark T. W. Ebbert

## Abstract

**Objective:** To systematically review and meta-analyze bulk RNA sequencing studies comparing Alzheimer’s disease (AD) patients with controls in human brain tissue, assessing study quality and identifying key genes and pathways.

**Methods:** We searched PubMed, Web of Science, and Scopus on September 23, 2023, for studies using bulk RNAseq on primary human brain tissue from AD patients and controls. Excluded were non-primary tissue, re-analyses without new data, limited RNA types and gene panels. Quality was assessed with a 10-category tool. Meta-analysis used high-quality datasets.

**Results:** From 3,266 records, 24 studies met criteria. Meta-analysis found 571 differentially expressed genes (DEGs) in temporal lobe and 189 in frontal lobe; overlapping pathways included "Tube morphogenesis" and "Neuroactive ligand-receptor interaction."

**Limitations:** Study heterogeneity and limited data tables constrained the review.

**Conclusions:** Rigorous methods are vital in AD transcriptomic studies. Findings enhance understanding of transcriptomic changes, aiding biomarker and therapeutic development.

**Registration:** PROSPERO (CRD42023466522).

## Introduction

Alzheimer’s disease (AD) is the most prevalent cause of dementia, affecting more than 33 million people worldwide, ultimately imposing over a trillion dollars of financial burden world-wide in 2019^1–4^. With the aging population, the number of individuals affected is projected to nearly triple, rising to over 90 million by 2050^1–4^. AD is characterized by the accumulation of amyloid beta plaques outside neurons and neurofibrillary tau tangles within neurons, along with neuronal death, which are hallmark features of the disease^5^. One key challenge in treating AD is that, by the time symptoms appear, substantial damage has already been done, limiting the effectiveness of treatments^6^. AD has a strong genetic component, with heritability estimated between 60% and 80%^7,8^. A recent genome-wide association study by Bellenguez et al. reported 75 loci associated with AD risk^9^, yet the mechanisms through which most of these loci affect disease development remain unclear. Shade et al. recently made an important step forward by beginning to clarify which AD-related genes are driving individual neuropathology endophenotypes, including discovering four new genes associated with AD and related dementias^10^, but much work still remains. Understanding the mechanisms driving AD is crucial for discovering presymptomatic biomarkers and improving treatments.

Transcriptomic studies in AD have been proposed to enhance our understanding of the disease’s mechanisms. Early research primarily focused on targeted approaches to examine genes associated with AD, where researchers identified four amyloid precursor protein (*APP*) mRNA isoforms and their potential roles in AD^11–14^. Similarly, targeted transcriptomic studies explored mRNA isoforms of the tau gene (*MAPT*) and their possible implications in the disease^15–19^. While targeted transcriptomic studies contributed important discoveries to AD research, their low throughput nature limited the number of genes that could be investigated. The second wave of transcriptomics studies in AD came with the advent of microarrays, which utilized probes to measure the expression of thousands of mRNA species at once, allowing researchers to compare their expression between brains of AD cases and controls. Microarray studies spanned multiple brain regions and revealed disrupted pathways in AD brains such as calcium signaling, inflammation, immune responses, apoptosis, oxidative stress, energy metabolism, and synaptic transmission^20–27^. Despite these improvements, microarrays can only measure a limited number of previously characterized RNAs. With the advent of short-read RNA sequencing (RNAseq), researchers could measure gene-level mRNA expression across the entire genome, including RNAs that were previously uncharacterized. Over the past decade, bulk tissue short-read RNAseq has become the standard method for comparing gene expression in diseased versus non-diseased tissues^28^. This approach has provided valuable insights into treatment targets and biomarkers for various human diseases.

For instance, RNAseq studies shed light on the interactions between the immune system and cancer, informing immunotherapy treatments^29^. Additionally, RNAseq characterized tumor subtypes and identified new potential drivers for medulloblastoma^30^, a highly malignant childhood brain tumor. For neuropsychiatric disorders, RNAseq has revealed new potential mechanisms and targets for conditions such as autism spectrum disorder, schizophrenia, and bipolar disorder^31^. Together, these findings demonstrate that leveraging bulk RNAseq is a valuable approach to studying complex diseases.

Similar to other disciplines, there is substantial research comparing gene expression in diseased versus non-diseased tissues in AD. Our objectives for this systematic review and meta-analysis are threefold: (1) to provide a comprehensive overview of bulk RNA sequencing studies conducted on human brain tissue, comparing AD patients with non-demented controls; (2) to assess the quality of these studies, identifying their strengths and weaknesses to guide the methodology for future transcriptomic research in AD; and (3) to conduct a meta-analysis with the most robust and extensive AD bulk RNA sequencing datasets available, offering insights into the genes and pathways that may be important for AD treatment and early diagnostics.

## Methods

### Search Strategy

A thorough search was conducted across PubMed, Web of Science, and Scopus on September 23^rd^, 2023, employing a comprehensive set of search terms related to transcriptomic changes in AD. The terms used were: (nanostring OR “RNA isoform” OR transcriptome OR transcriptomic OR RNA-seq OR RNAseq OR "RNA microarray" OR "mRNA microarray" OR "RNA sequencing") AND (Alzheimer OR Alzheimer’s) AND (human OR man OR woman OR men OR women OR patient OR patients OR "homo sapiens"). Duplicates were removed within each database, and the results were merged using Mendeley (https://www.mendeley.com/) where duplicates were again identified and removed. The records were then imported into Rayyan (https://www.rayyan.ai/), a platform for systematic review management, where duplicates were further identified and removed.

### Study Selection

Each record’s title and abstract were screened in Rayyan by at least two team members from a six-member team working independently. Decision conflicts were resolved through discussion. Records were evaluated based on study design, study population, and outcome measures. Our inclusion criteria were as follows: (1) AD compared to a control group; (2) early or late-onset AD; (3) original data or reanalysis of previous data; (4) RNAseq, cDNA/RNA microarray, or Nanostring techniques; and (5) transcriptomic data on primary human tissue (not a cell line or microbiome sample). Exclusion criteria were: (1) other review articles, case reports, book chapters, posters, editorials; (2) not a transcriptomics study; (3) RT-qPCR as the only transcriptomic approach; and (4) transcriptomic analyses on non-primary human tissue such as mouse brain or human cell line.

Records meeting the inclusion criteria proceeded to the next step. The inclusion criteria for the title and abstract screening were intentionally less stringent than those for the full-text assessment to avoid excluding relevant records due to incomplete information in the title and abstract.

Next, a detailed full-text assessment was conducted using refined inclusion criteria: (1) AD compared to non-demented controls; (2) late-onset AD cases; (3) primary human brain tissue samples; (4) presenting original data; (5) bulk RNAseq techniques; (6) traditional bulk RNAseq measurements (mRNA, long non-coding RNA, polyA RNA, depleted ribosomal RNA); and (7) differential gene and/or RNA isoform expression analysis.

Exclusion criteria were: (1) other review articles, case reports, book chapters, posters, editorials; (2) no primary human brain tissue; (3) reanalysis of publicly available data without presenting any original RNAseq data fitting inclusion criteria; (4) no bulk RNAseq data; (5) studies limited to small and/or circular RNAs; (6) no comparison between AD subjects and non-demented controls; (7) no differential gene or RNA isoform expression analysis (8) analysis limited to a subset/panel of target genes; and (9) main text not in English.

One author thoroughly scrutinized each article, identifying 24 eligible studies, as depicted in **Figure 1**. We made one exception during the full-text review: although the record by Marques-Coelho et al.^32^ did not include original data, it was deemed eligible because it was the only study we found that conducted both differential gene expression and differential isoform usage analyses using the largest publicly available bulk RNAseq cohorts from the AD Knowledge Portal^33^ (https://adknowledgeportal.synapse.org/). They separately analyzed data from the Mayo Clinic cohort^34^, the Mount Sinai/JJ Peters VA Medical Center Brain Bank (MSBB) cohort^35^, and the Religious Orders Study and Rush Memory and Aging Project (ROSMAP) cohort^36^. Basic information for each study can be found in **Table 1**.

**Figure 1:**
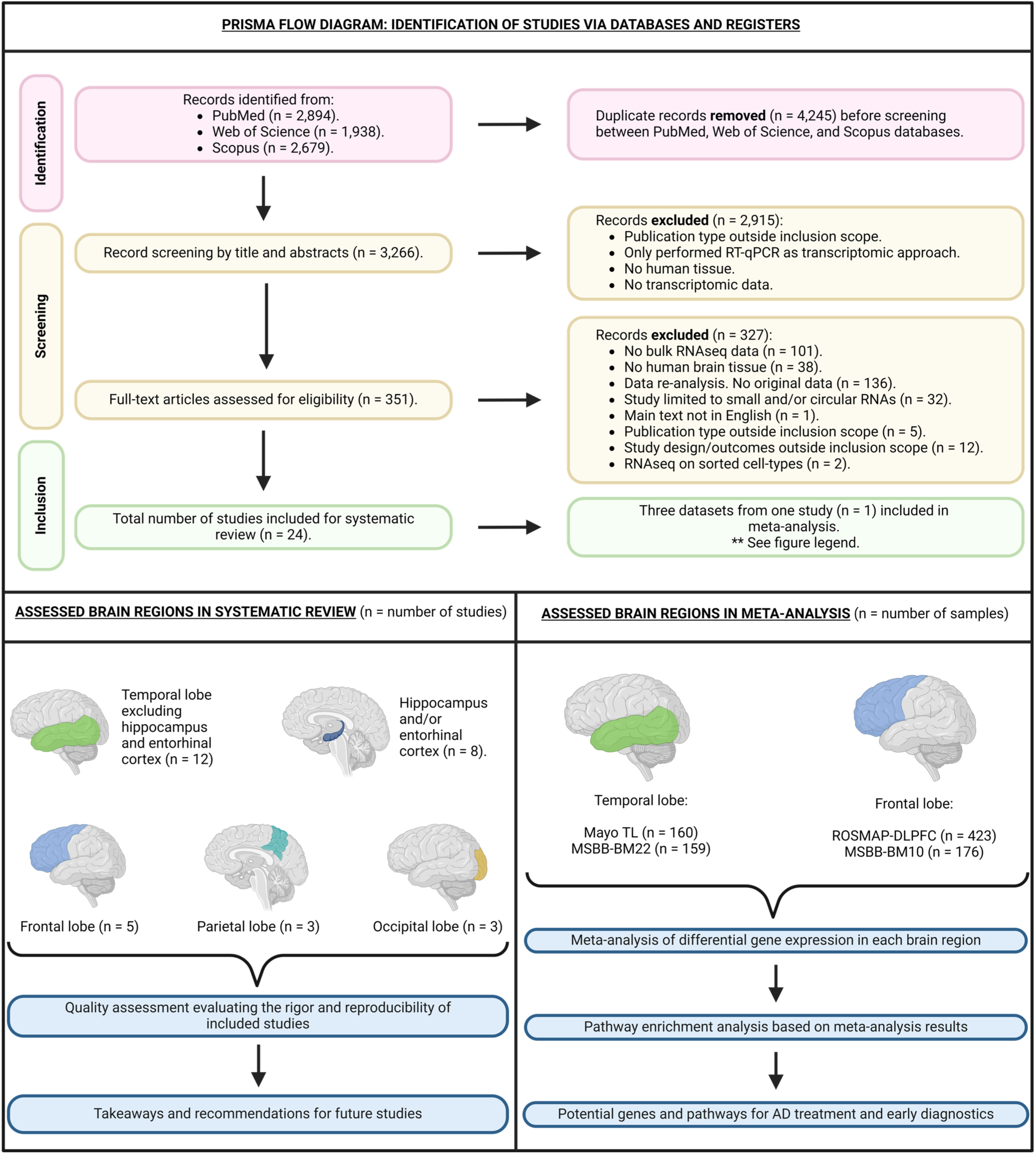
PRISMA flow diagram and study design. Diagram illustrating the process to identify studies that met inclusion criteria. The diagram also shows the assessed brain regions in the systematic review and the corresponding number of studies for each region. The total number of studies across all brain regions exceeds the total number of included studies (n = 24) because some studies evaluated multiple brain regions. **Only three datasets from one study were included in the meta-analysis for the following reasons: (1) the study had a significantly larger sample size compared to others; (2) it received the highest quality assessment score, reducing the risk of bias; and (3) only five of the remaining 23 studies provided a differential expression table suitable for meta-analysis, all of which had relatively small sample sizes (n < 50) or significant methodological issues, such as poorly defined AD status.

**Table 1:** Brief overview of study characteristics. Table showing first author and year for each study, brain regions assessed, and sample size per group. More thorough overview of study characteristics provided in the data extraction table (Supplementary Table 2). A “(?)” in the “Sample size per group” column indicates that the study omitted information about number of females included. Table is organized by brain region and then sorted by year of publication.

| Study (year of publication) | Brain region(s) | Sample size per group: total number (number of females) |
| --- | --- | --- |
| **Temporal lobe (excluding hippocampus & entorhinal cortex)** |  |  |
| Humphries et al. (2015) | Temporal pole | AD: 10 (0) \| Control: 10 (0) \| Dementia with Lewy Bodies: 10 (0) |
| Barbash et al. (2017) | Temporal lobe | RUSH cohort: Non-demented 4 (0) \| MCI & Low Braak 4 (0) \| MCI and Medium Braak 4 (0) \| MCI & High Braak 4 (0) \| Demented & Low Braak 4 (0) \| Demented & Medium Braak 4 (0) \| Demented & High Braak 4 (0)<br>NBB cohort: Healthy controls 8 (0) \| Early pathology but non-demented 8 (0) \| Advanced AD pathology & cognitive decline 8 (0) |
| Piras et al. (2019) | Middle temporal gyrus | AD: 8 (5) \| Control: 8 (4) |
| Cho et al. (2020) | Anterior temporal lobe | Chronic traumatic encephalopathy (CTE): 8 (0) \| CTE/AD: 6 (0) \| AD: 10 (6) \| Control: 10 (4) |
| Felsky et al. (2022) | Temporal cortex | Caribbean Hispanic AD: 23 (16) \| Caribbean Hispanic Control: 16 (8) |
| Das et al. (2023) | Superior temporal gyrus | AD: 10 (5) \| Control: 8 (4) |
| **Hippocampus & entorhinal cortex** |  |  |
| Magistry et al. (2015) | Hippocampus | AD: 4 (3) \| Control: 4 (2) |
| Annese et al. (2018) | Hippocampus | Control: 5 (0) \| AD: 5 (0) \| Parkinson's disease: 6 (0) |
| van Rooij et al. (2019) | Hippocampus | AD = 18 (10) \| Control: 10 (5) |
| Jia et al. (2021) | Entorhinal Cortex | AD: 7 (4) \| Control: 18 (7) |
| Shmookler Reis et al. (2021) | Hippocampus | AD: 3 (?) \| Control: 3 (?) |
| Luo et al. (2023) | Hippocampus & Entorhinal Cortex | AD: 31 (16) \| Control: 22 (8) |
| **Frontal lobe** |  |  |
| Panich et al. (2021) | Dorsolateral pre-frontal cortex. | Framingham Heart Study & Boston University Alzheimer's Disease Center<br>AD: 64 (27) \| Controls: 129 (65) |
| Fisher et al. (2023) | Prefrontal cortex | Control: 16 (?) \| AD: 13 (?) \| Lewy body dementia: 12 (?) \| Lewy body dementia & AD: 10 (?) \| Parkinson's disease = 8 (?) |
| **Parietal & occipital lobe** |  |  |
| Mills et al. (2013) | Parietal Lobe | AD: 5 (0) \| Control: 5 (0) |
| Guennewig et al. (2021) | Primary visual cortex & precuneus | AD: 5 (2) \| Control: 5 (2) |
| Caldwell et al. (2022) | Primary visual cortex | Early onset sporadic AD: 19 (7) \| Late-onset AD: 20 (9) \| Control: 8 (4) |
| **Multiple brain regions** |  |  |
| Twine et al. (2011) | Total brain, frontal lobe, & temporal lobe | Total brain control = 23 (10) \| Frontal lobe control = 10 (5) \| Total brain AD = 1 (0) \| Frontal AD = 1 (0) \| Temporal lobe AD = 1 (0) |
| Miller et al. (2017) | Temporal cortex, parietal cortex, parietal white matter, & hippocampus. | Non-demented = 56 (20) \| Demented = 50 (23) |
| Lee et al. (2020) | Cortex tissues | Alzheimer's disease: 6 (?) \| Controls: 6 (?) |
| Li et al. (2021) | Superior temporal gyrus & inferior frontal gyrus | Superior temporal gyrus AD = 24 (11) \| Mild Cognitive Impairment (MCI) = 14 (7) \| Control = 38 (11).<br>Inferior frontal gyrus - AD: 33 (21) \| MCI: = 9 (4) \| Control = 23 (6). |
| Marques-Coelho et al. (2021) | Frontal lobe & temporal Lobe | Mayo Temporal Lobe - AD: 82 (49) \| Control: 78 (37)<br>Mount Sinai Brodmann Area 10 - AD: 105 (65) \| Control: 71 (39)<br>Mount Sinai Brodmann Area 22 - AD: 98 (57) \| Control: 61 (32)<br>Mount Sinai Brodmann Area 36 - AD: 88 (56) \| Control: 64 (33)<br>Mount Sinai Brodmann Area 44 - AD: 90 (56) \| Control: 63 (31)<br>ROSMAP Dorsolateral Prefrontal Cortex - AD: 222 (155) \| Control: 201 (124) |
| King et al. (2022) | Primary visual cortex & middle temporal gyrus | Mid-life control: 15 (5) \| Healthy aged control: 18 (7) \| AD = 13 (6)<br>Lifetime cognitive resilient = 9 (4) \| Lifetime cognitive decline = 8 (2) |
| Santana et al. (2022) | Auditory cortex, hippocampus, & cerebellum. | AD: 6 (4) \| Control: 6 (3) |

### Data Extraction

Two authors working independently extracted relevant data from the 24 included studies. Data was gathered for the following categories: “study groups included”, “brain region sampled”, “sex of samples”, “sample size per group”, “sample size per race/ethnicity”, “reported age per group”, “Alzheimer’s disease diagnostic criteria”, “years of AD diagnosis”, “RNA extraction kit”, “RNA enrichment or depletion”, “RNA integrity score”, “Library preparation”, “post-mortem interval”, “read length”, “depth of sequencing per sample”, “paired or single end reads”, “sequencing platform”, “aligner”, “reference genome”, “quantification tool”, “transcriptome annotation”, “transcript discovery method”, “differential expression tools”, “covariates in differential expression analysis”, “expression filtering”, “multiple testing correction”, “differential expression thresholds”, “differentially expressed genes results”, and “table provided for all genes included in differential expression analysis”. A definition for all data extraction terms is provided in **Supplementary Table 1**. Data that was not available was listed as ‘Not Reported’. Any assumptions made about data are reported in the data extraction table (**Supplementary Table 2**).

### Methodological Quality Assessment

We evaluated each study’s methodological quality, which also serves as a proxy for risk of bias in this study. Two authors working independently scored each study and conflicts were resolved through discussion.

Factors were assessed on a three-point scale for: “sample size” “sex and ethnicity”, “AD diagnostic criteria”, “control matching”, “transcript level analysis”, “results validation”, “sequencing depth”, “statistical rigor”, “data availability”, and “reproducibility”. The objective criteria used for scoring each category are described in **Table 2**. The points for all categories were averaged to give the overall quality assessment score to each study. We tested for correlations between year of publication and each quality assessment category using Spearman’s rho. Correlations were considered statistically significant if they had a Bonferroni corrected p-value < 0.1 (equivalent to unadjusted p-value < 0.009). We used custom Python scripts to perform this analysis and generate visualizations (**Code Availability**).

**Table 2:**
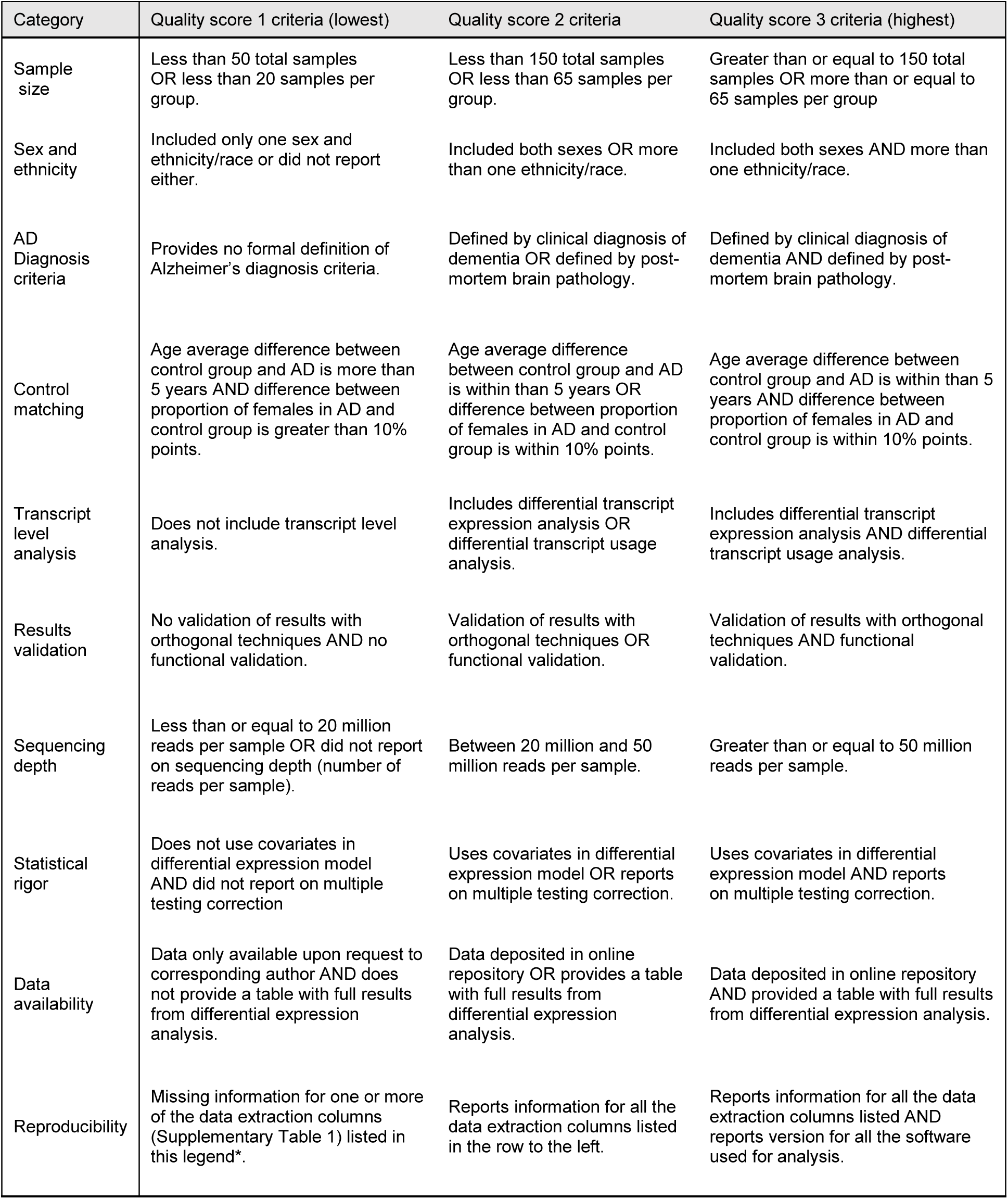
Definitions of objective criteria for quality assessment scoring. The lowest/worst quality score is one (1) and the highest/best quality score is three (3). There are 10 different categories for objective quality assessment. The data extraction table (**Supplementary Table 2**) columns used to score the “reproducibility” category are: RNA extraction kit, RNA enrichment and/or depletion, RNA Integrity Score (RIN), Library preparation, Post-Mortem Interval (PMI), Read Length, Number of Reads per sample, Paired-end or single end reads, Sequencing platform, Aligner, Reference genome, Quantification tool, Transcriptome annotation, Differential expression tools, Differential expression thresholds.

### Meta-analysis

We conducted a meta-analysis using three datasets examined by Marques-Coelho and colleagues^32^, namely Mayo clinic temporal lobe^34^, MSBB frontal and temporal lobes^35^, and ROSMAP dorsolateral prefrontal cortex^36^. Three datasets from this single study were selected for the meta-analysis due to the following factors: (1) the study had a considerably larger sample size than all others; (2) it achieved the highest quality assessment score, minimizing the risk of bias; and (3) only five out of the remaining 23 studies provided a differential expression table appropriate for meta-analysis, all of which had either small sample sizes (n < 50) or substantial methodological concerns, such as loosely defined AD status. We divided the datasets analyzed by Marques-Coelho et al.^32^ into two separate meta-analyses. The first focused on the temporal lobe, using MSBB Brodmann area 22 (n = 159) and Mayo temporal lobe data from Brodmann areas 20/21/22/41/42 (n = 160).

The second meta-analysis centered on the frontal lobe, utilizing MSBB Brodmann area 10 (n = 176) and ROSMAP Brodmann areas 9/46 (n = 423). Two MSBB datasets analyzed by Marques-Coelho et al^32^, MSBB Brodmann area 44 (frontal lobe) and MSBB Brodmann area 36 (temporal lobe), were excluded from our analysis due to overlapping subjects with the included datasets, which would violate the critical assumption that data must be independent meta-analyses. MSBB Brodmann area 22 was chosen for the temporal lobe analysis over Brodmann area 36 due to its slightly larger sample size and greater anatomical and functional similarity to the Mayo temporal lobe dataset brain regions. For the frontal lobe analysis, MSBB Brodmann area 10 was selected over Brodmann area 44 for its larger sample size and closer anatomical and functional resemblance to the ROSMAP dataset’s Brodmann areas.

The original publication did not provide tables suitable for meta-analysis because the journal would not publish such a large file (215 megabytes). We contacted the corresponding author to obtain a table containing p-values and log2FoldChange values for all genes analyzed in each dataset. The corresponding author provided the table with adjusted p-values, which was available with their pre-print submission for the article. The unadjusted p-values were not available; therefore, we performed the meta-analysis using the Benjamini-Hochberg adjusted p-values provided by the corresponding author—ultimately making our meta-analysis more stringent and conservative.

We restructured the tables using a custom Python script to ensure the tables met input criteria for METAL^37^, which we used to perform the meta-analyses. We also excluded genes that were not common to all datasets within a brain lobe (**Code Availability**). Statistics for the meta-analysis are derived from the inverse-variance weighted fixed-effect model from METAL. We set the weight of each dataset to its sample size and applied a Bonferroni correction to the p-value outputs from our meta-analysis. Genes were considered differentially expressed between AD subjects and non-demented controls if the Bonferroni adjusted p-value was < 0.1. We included 29,492 genes in the temporal lobe meta-analysis, resulting in an unadjusted p-value threshold of 3.39 × 10⁻^6^. For the frontal lobe meta-analysis, we included 31,378 genes, resulting in an unadjusted p-value threshold of 3.19 × 10⁻^6^. The I^2^ metric for heterogeneity, calculated using METAL, was moderate to low. We chose not to explore the causes of heterogeneity due to the small number of datasets, which would make identifying the probable causes difficult. Results for the differential expression meta-analysis were tabulated and visually displayed using a custom Python script (**Code Availability**).

We then performed a pathway analysis for up- and down-regulated differentially expressed genes using Metascape^38^ (https://metascape.org) to identify potentially disrupted pathways in the brains of AD patients. The Metascape analysis was performed separately for the temporal lobe and frontal lobe. Figures depicting pathway analysis results were generated by Metascape.

### Overlap Analysis

We examined the overlap between the differentially expressed genes (DEGs) from our meta-analysis and those reported in the studies included in this systematic review for the temporal and frontal lobes. Eight temporal lobe studies provided differential gene expression tables suitable for this comparison, while three studies did so for the frontal lobe. For each DEG identified in the temporal lobe meta-analysis, we tallied how many of the other temporal lobe studies reported the same DEG in the same direction (e.g., upregulated in AD) We applied the same process to the frontal lobe data. DEGs that did not appear in any other study were considered unique to our meta-analysis.

### Rigor and Reproducibility

A Singularity container was used for all analyses in the present study. Singularity containers bundle software components in a portable and reproducible manner^39^. All the scripts in this study and the instructions on how to access the singularity container used to execute them can be found in the GitHub repository for this project (**Code Availability**). The versions for all software used in the analysis can also be found on the project’s GitHub under the singularity container definition file and in **Supplementary Table 3.** Additional files such as meta-analysis input files, meta-analysis output files, and Metascape output were deposited in an open Zenodo repository (**Data Availability**). This systematic review and meta-analysis was conducted following PRISMA guidelines^40^, see checklists in **Supplementary Files 1 and 2**. This review was registered on PROSPERO, registration number: CRD42023466522. The original review protocol can be accessed here: https://www.crd.york.ac.uk/prospero/display_record.php?RecordID=466522

## Results

The initial literature search yielded 7,528 records, ultimately resulting in 3,266 unique records after merging and removing duplicates. Following title and abstract screening, 2,915 records did not meet inclusion criteria, leaving 351 records for full-text screening. Upon thorough examination of the full texts, 24 studies conducting bulk RNAseq on AD brain tissue met the eligibility criteria and were included in the review (**Figure 1**). One notable study that was close to fitting the inclusion criteria, but was excluded from our systematic review was Raj et al.^41^. The article analyzes ROSMAP data, but the outcomes are mainly focused on differential intron usage, splicing quantitative trait loci analysis, and transcriptome wide associations rather than differential gene and/or transcript expression analysis.

### Study Characteristics

The dataset included publications spanning from 2011 to 2023. Among these, two records had sample sizes exceeding 150,^32,42^ three had sample sizes ranging from 50 to 150,^43–45^ and 19 had fewer than 50^46–64^ (**Table 1**). The temporal lobe excluding hippocampus and entorhinal cortex was the most frequently sampled brain region, representing 12 studies.^32,43,45,46,51,52,55,56,60–62,64^. Sampling specific to the hippocampus or entorhinal cortex was performed in eight studies.^44,45,48–51,53,59^ Other regions studied included the frontal lobe (five studies),^32,42,43,46,58^ the parietal lobe (three studies),^45,47,57^ and the occipital lobe (three studies).^55,57,63^ One record did not specify the brain lobe studied, describing the brain region as “cortex tissues”.^54^

In addition to AD cases and non-demented controls, two records included a neurodegenerative disease control group,^56,58^ and two papers examined cohorts with AD occurring concurrently with another neurologic disease, such as dementia with Lewy bodies.^58,62^ All records performed differential gene expression analysis,^32,42–64^ with three also conducting differential transcript expression or differential transcript usage analysis.^32,46,47^

Regarding population representativeness, 17 records included both men and women,^32,42–45,48,50–53,55,57,58,60–63^ five studies included only men,^46,47,56,59,64^ and two studies did not specify the gender distribution within their cohorts.^49,54^ Multiple ethnicities were only present in one study.^61^ Six studies focused on cohorts from a single ethnicity^44,45,48,56,58,59^—Chinese or Caucasian—while ethnicity was not reported in seventeen studies.^32,42,43,46,47,49–55,57,60,62–64^

Below is a summary of the included papers, organized by the brain region studied. If a paper investigates multiple regions but contains original data for only one, it is discussed in the section relevant to the brain region where the original data stems from. Detailed information about data extraction definitions and the complete data extraction table are provided in **Supplementary Tables 1 and 2**.

### Studies in temporal lobe excluding hippocampus and entorhinal cortex

There were six studies in temporal lobe specifically (excluding studies specifically on the hippocampus and entorhinal cortex); here, we briefly summarize each study in order of publication. Humphries et al.^56^ (published in 2015) included patients with AD, cognitively normal controls, and a disease control group (dementia with Lewy Bodies). Including this disease control group allowed for differentiation between processes due to general neurodegeneration and those specific to AD. Each group had 10 subjects, with one temporal lobe sample per subject. The study identified 16 differentially expressed genes (DEGs) when comparing AD patients to normal controls. Further comparison of these genes between AD and the disease control group revealed that five genes were differentially expressed—*C10orf105* and *RARRES3* were upregulated in AD while *DIO2*, *ENSG00000249343.1*, and *WIF1* were downregulated. These transcriptional findings were validated using two independent, publicly available microarray datasets. Additionally, network analysis of 2,504 genes with nominal transcription differences between AD and normal controls indicated differences in myelination and innate immunity. Study strengths include well-defined AD diagnostic criteria through cognitive and pathological assessments, the inclusion of a neurodegenerative disease control group, validation of results with independent datasets, and high sequencing throughput (over 50 million reads per sample). However, the study’s limitations are a small sample size (10 subjects per group), a lack of females, not sharing raw data in an online repository, the omission of a comprehensive table of differential gene expression analysis results, and not reporting on key methodological details such as post-mortem interval.

Barbash et al.^64^ (published in 2017) used temporal lobe samples from the RUSH Memory and Aging project, divided into cognitively normal controls, mild cognitive impairment, and patients with AD. Each group was further categorized by low, moderate, and high Braak staging, resulting in nine groups with four subjects each. Additionally, temporal gyrus samples obtained from the NeuroBioBank were divided into patients with AD, non-demented patients with pathologic AD, and non-demented patients without pathologic AD, with eight samples per group. These samples underwent RNAseq after selective quantitative amplification of RNA to identify transcript variants with alternative 3’-UTR alternatively poly-adenylated sequences. They reported 1217 DEGs associated with cognition and 570 DEGs associated with pathology. Differential polyadenylation analysis showed 98 genes with increased adenylation in AD and non-demented with pathologic AD compared to non-demented without pathologic AD. In addition, there were 45 genes with decreased adenylation in AD and non-demented patients with pathologic AD compared to non-demented patients without pathologic AD. Lipid processing, cognition level, and AD pathology were linked, implicating genes *NOVA1* and *hnRNPA1* in patients with AD pathology but normal cognition near the time of death. Strengths of this study include well-defined AD diagnostic criteria through cognitive and pathological assessments, subgroup analysis stratified by cognitive and pathological criteria, results validation with lipid mass spectrometry and RT-PCR, assessment of cell-type specific gene expression, RNAseq data deposited in an online repository, and high sequencing throughput (over 50 million reads per sample). However, the study’s limitations are a small sample size, a lack of females, the omission of a comprehensive table of differential gene expression analysis results, and missing key methodological details such as post-mortem interval for RUSH Memory and Aging project samples.

Piras et al.^52^ (published in 2019) performed RNAseq of the middle temporal gyrus in eight AD patients and eight non-demented controls. They identified 1,534 differentially expressed genes. Upon comparison with independent datasets, 453 validated genes were found. Pathway analysis of these validated genes revealed Clathrin-mediated endocytosis as the most significantly associated pathway. Further analysis correlated the validated genes with Braak staging and neurofibrillary tangles, identifying *AEBP1* (upregulated) and *NRN1* (downregulated) as the most significantly linked genes and potential biomarkers for AD. Additionally, pathway analysis specific to long non-coding RNAs revealed disruptions in 61 pathways, including those involved in GABAergic transmission and peptide chain elongation. The study’s limitations include a small sample size, the use of short-read RNAseq (50 nucleotides) for transcriptome assembly which may lead to inaccuracies, the absence of raw data in an online repository, lack of a comprehensive table with differential gene expression analysis results, and not reporting participants’ ethnicity. The study strengths included well-defined AD diagnostic criteria through cognitive and pathological assessments, a thorough description of methodology enhancing reproducibility, inclusion of both sexes, and validation of main findings using an independent microarray dataset.

Cho et al.^62^ (published in 2020) focused on comparing patients with chronic traumatic encephalopathy (CTE), AD, CTE concurrent with AD (CTE/AD), and normal controls. RNAseq was performed on anterior temporal lobe samples to investigate transcriptomic differences. Using weighted gene co-expression network analysis (WGCNA) and principal component analysis (PCA), the study found that genes related to synapse signaling, such as synaptotagmins, were commonly downregulated in CTE, CTE/AD, and AD. Genes involved in memory function, including calcium/calmodulin-dependent protein kinase II, protein kinase A, protein kinase C, and AMPA receptor genes, were downregulated across these disorders, highlighting similarities between AD and CTE. These RNAseq findings were validated using qPCR and Western blot analyses. Strengths of this study include availability of raw data in an online repository, validation of results using orthogonal methods, and the inclusion of CTE and CTE/AD groups. Limitations of this study include lack of well-defined AD diagnostic criteria, small sample size, inclusion of females in only two out of the four groups, omission of a comprehensive table with differential gene expression analysis results, and omission of key details such as post-mortem interval and RNA integrity number.

Felsky et al.^61^ (published in 2022) was the only literature record meeting our inclusion criteria that investigated AD transcriptomic differences among different ethnic groups, making it a particularly unique and important study. It performed RNAseq on the temporal cortex of Caribbean Hispanic patients and controls, comparing results to non-Hispanic white cohorts in the ROSMAP and Mayo datasets. They found 118 genes with significant effects in Caribbean Hispanic patients, showing opposite directions in both comparison datasets, suggesting ancestry-driven differences in translational machinery activation. *NPNT* was the top differentially expressed gene in the cross-ancestry meta-analysis. Ancestry-specific enrichment analysis highlighted ribosomal genes and those involved in protein synthesis and trafficking. The strengths of this study include the representation of two ethnicities and both sexes, validation and comparison of results using publicly available RNAseq data, and providing a comprehensive table with differential gene expression analysis results. The study is limited by the relatively low RNA quality in the Caribbean Hispanic cohort, small sample size, lack of raw data availability for newly generated data, and missing key methodological details such as not specifying which genome annotation was used.

Das et al.^60^ (published in 2023) utilized laser capture microdissection on superior temporal gyrus samples from AD patients and controls. In AD patients, areas dissected included amyloid beta plaques, the halo surrounding these plaques, neurofibrillary tangles (NFT) and their halo, and regions free from plaques and tangles. These were compared to plaque and tangle free cortical areas in normal controls. Additionally, the AD patients were subdivided based on APOE status into ε3/ε3 and ε4/ε4 groups to examine within-group differences. The study found that amyloid beta plaques showed an upregulation of microglial genes and a downregulation of neuronal genes, indicating neuroinflammation and phagocytosis. In contrast, NFTs were only associated with downregulation in neuronal genes. When comparing the APOE ε3/ε3 and ε4/ε4 groups, the transcriptomic changes were more pronounced in the amyloid beta plaques of the ε4/ε4 group. The strengths of this study include an innovative approach using laser capture microdissection to understand transcriptomic changes associated with neurofibrillary tangles and beta-amyloid plaques. Additionally, it features well-defined AD diagnostic criteria through cognitive and pathological assessments, subgroup analysis based on APOE status, a comprehensive table of differential gene expression analysis results, and deposition of raw data in an online repository. However, the study is limited by a small sample size, lack of validation for results, and the use of short reads (60 nucleotides) for transcriptome assembly which may lead to inaccuracies.

### Studies in hippocampus and entorhinal cortex

Six studies focused specifically on the hippocampus and entorhinal cortex, where AD pathogenesis generally begins. Magistri et al.^53^ (published in 2015) compared RNAseq differences in the hippocampus of four late-onset AD cases versus four age-matched controls. Differential expression analysis identified 143 protein-coding genes, 90 lincRNAs, 31 antisense RNAs, and 1 novel putative protein-coding gene with significant differences between AD and controls. Of these, 61 protein-coding genes were under-expressed and 82 were over-expressed in AD. Notably, the *TAC1* gene, encoding substance P, was downregulated, and *SERPINE1*, coding for plasminogen activator inhibitor type-1, was significantly upregulated in AD. Pathway analysis highlighted neurovascular defects and altered amyloid-β homeostasis. Additionally, 72 of the differentially expressed lincRNAs were novel, with AD-linc1 and AD-linc2 validated by RT-qPCR. Among differentially expressed antisense RNAs, 21 were upregulated and 10 downregulated. Strengths of this study include RT-qPCR and cell-line functional validation of results, well-defined AD diagnostic criteria through cognitive and pathological assessments, high Braak scores in AD samples without cellular population imbalances compared to controls, high sequencing depth (over 50 million reads per sample), and sharing raw data in an online repository. However, the study has some limitations including small sample size, missing key methodological details such as read length, and not providing a table with comprehensive differential gene expression analysis results.

Annese et al.^59^ (published in 2018) compared RNAseq results from the hippocampal CA1 region of six late-onset AD patients, six cognitively normal controls, and six Parkinson’s disease patients as disease controls, including both grey and white matter. Principal component analysis identified and excluded one control and one AD sample as outliers. AD cases were selected based on dementia status, Braak V or VI, and positive Aβ plaques and NFTs on autopsy, while controls had no neurological disease history or brain abnormalities.

Parkinson’s disease patients were positive for Lewy bodies in the substantia nigra. The study identified 2,122 DEGs between AD and controls, with 2,075 protein-coding genes (789 upregulated, 1,286 downregulated) and 47 lncRNAs (19 upregulated, 28 downregulated). Between Parkinson’s disease samples and non-demented controls there were 19 differentially expressed protein-coding genes, with 11 overlapping with AD, leaving 2,064 genes for pathway analysis. Additionally, small RNAseq was conducted on the hippocampus, middle temporal gyrus, and middle frontal gyrus in AD patients and controls, and only the hippocampus in Parkinson’s disease patients. RNAseq results were validated by RT-qPCR in a larger cohort of nine controls and nine AD patients. The study confirmed no significant changes in neuronal marker expression, indicating no cell population imbalances. Strengths of this study include RT-qPCR validation, thorough cell population assessment, well-defined AD diagnostic criteria through cognitive and pathological assessments, high sequencing depth (over 50 million reads per sample), and exemplary reporting of methods, including all software versions used and major methodological details. Additionally, the study included a disease control (Parkinson’s disease). The main limitations are the small sample size, lack of raw data sharing in an online repository, absence of a comprehensive table of differential gene expression analysis results, and not including females.

The study by van Rooij et al.^50^ (published in 2019) compared whole transcriptome sequencing of 18 AD hippocampal samples to 10 age- and sex-matched cognitively healthy controls. Two outlier AD cases were removed due to high expression of *TTR*, a gene specifically expressed in the choroid plexus. The study did not provide specific parameters for defining AD cases versus controls, but post-mortem analysis revealed significant differences in Braak score, brain pH, brain weight, amyloid deposition, and post-mortem delay, with Braak scores above five for AD cases and below 2.8 for controls. RNA was isolated from the dentate gyrus and cornu ammonis for each case and control. The researchers identified 2,716 DEGs in the discovery dataset, with 1,610 DEGs involved in protein-protein interactions, clustering 735 DEGs into 33 gene modules. These findings were replicated in an independent RNAseq dataset, identifying 2,490 DEGs with 1,311 replicating from the discovery set, clustering 653 DEGs into 37 modules. Gene set enrichment analysis revealed significant gene ontology biological processes in each module. The study’s strengths include the replication of findings using a publicly available RNAseq dataset, and the provision of a comprehensive table of differential gene expression analysis results. However, there are several limitations. The study does not report key details such as RNA integrity scores and the ethnicity of subjects, nor does it provide formal criteria for AD diagnosis.

### Additionally, the sample size is relatively small

Jia et al.^48^ (published in 2021) took postmortem entorhinal cortex samples from AD patients and non-AD controls to investigate both proteomic and transcriptomic changes in the disease. Proteomic analysis, performed with liquid chromatography tandem mass spectrometry on tissues from four AD and four non-AD donors, identified differentially expressed proteins, while transcriptomic analysis, using RNA sequencing on seven AD and 18 control samples, identified differentially expressed genes. Integrated transcriptomics and proteomic analysis revealed substantial dysregulation of ion transport, further validated through immunohistochemistry. This study is notable for including both sexes and combining transcriptomic and proteomic data. However, it was limited by a small sample size, lack of detailed reporting on sequencing methods, insufficient data on cohort characteristics, and lack of publicly available raw data, impacting the study’s rigor and reproducibility.

Shmookler Reis et al.^49^ (published in 2021) explores the role of nucleic acids in protein aggregation linked to neurodegenerative diseases. The researchers extracted and sequenced RNA and DNA from Sarkosyl-insoluble aggregates taken from the hippocampus of three AD patients and three age-matched controls.

RNAseq revealed a significant, nonrandom presence of specific RNA sequences within the aggregates, particularly overrepresented in AD samples. Knockdown of the translational elongation factor *EEF2* in a cell-line model significantly reduced RNA content in the aggregates, underscoring the role of translational dynamics on aggregate composition. This innovative study explores the molecular pathology of neurodegenerative diseases, highlighting translational processes and nucleic acid interactions as potential therapeutic targets.

The study’s strengths include functional experimentation and epigenetic comparisons. However, it faces several limitations. These include a small sample size, not correcting p-values for multiple comparisons, lack of a comprehensive table of differential gene expression analysis results, failure to deposit raw data in an online repository, and insufficient methodological details. Specifically, the study does not provide a formal definition for AD, lacks information on sample demographics, RNA quality metrics, and specific RNA sequencing parameters, which significantly compromise the study’s rigor and reproducibility.

Luo et al.^44^ (published in 2023) conducted a thorough RNAseq analysis, incorporating cell-type deconvolution, surrogate variable analysis for batch effects, and WGCNA network analysis in a Chinese population. It examined five critical brain regions associated with AD: hippocampus CA1-CA4 and the entorhinal cortex. Key discoveries included identifying extensive gene expression changes, highlighting the role of *PSAP* in promoting astrogliosis and A1-reactive astrocyte phenotype, and uncovering AD-related signaling pathways and cell-type specific responses, particularly in the entorhinal cortex. The study’s strengths include its use of multiple brain regions, inclusion of both sexes, and functional follow-up involving animal models and in-vitro astrocyte-neuron co-culture from rat-derived cells. However, the study had limitations such as a relatively small sample size, vague methodological reporting (e.g., unspecified p-value correction, sequencing instrument, read length, and read type), and lack of raw data availability.

### Studies in frontal lobe

Of the 24 literature records meeting our inclusion criteria, only two performed RNAseq on the frontal lobe specifically. Panitch et al.^42^ (published in 2021) grouped patients with and without AD based on their *APOE* genotype, including ε2/ ε3, ε3/ ε3, and ε3/ ε4 groups. RNAseq was performed on samples from the prefrontal cortex, revealing differential expression of complement pathway genes, particularly *C4A* and *C4B* in *APOE* ε2/ ε3 AD cases compared to controls. These genes are involved in immune and inflammatory responses in the brain, suggesting a mechanism that may contribute to the protective effects of the ε2 allele against AD. The study also identified a specific co-expression network related to astrocytes, oligodendrocytes, and their progenitor cells, which is enriched in *APOE* ε2/ε3 AD cases. Key genes in this network, including *C4A*, *C4B*, and *HSPA2*, are significantly associated with tau pathology, indicating a tau-centric mechanism of AD progression in the context of *APOE* ε2. This study’s strengths include a large sample size, the inclusion of both sexes, and inclusion of publicly available datasets to supplement original data. Differential gene expression analysis was performed by *APOE* genotype, using well-defined AD diagnostic criteria through cognitive and pathological assessments. Additionally, well-matched control subjects were included, with similar age and sex distributions to the AD subjects. However, the study did not report participants’ ethnicity or share data in an online repository. It also lacked a comprehensive table of differential gene expression analysis results and omitted key details such as sequencing depth and read length for RNA sequencing data.

Fisher et al.^58^ (published in 2023) investigated differentially expressed and methylated genes in the prefrontal cortex across five groups: individuals with AD, pure dementia with Lewy bodies, dementia with Lewy bodies with concomitant AD, Parkinson’s disease, and non-demented controls. The researchers identified distinct methylation and transcriptional patterns between Lewy body driven dementias versus amyloid-beta driven dementias. Significantly, they discovered transcriptional differences in oxidative stress response pathways between controls and all dementia types, highlighting the role of these processes in dementia development and progression. The study’s strengths include the inclusion of multiple disease control groups as well as a non-demented control group, well-defined AD diagnostic criteria through cognitive and pathological assessments, shared raw data in an online repository, a comprehensive table of differential gene expression analysis results, and the inclusion of methylation data. However, the study’s limitations include a small sample size and not reporting key methodological details such as sequencing depth, read length, and whether reads were single-end or paired-end.

### Studies in parietal and occipital lobe

Three out of the 24 literature records performed bulk RNAseq in the parietal or occipital lobes specifically. Mills et al.^47^ (published in 2013) utilized RNAseq to investigate gene and transcript expression changes in the parietal cortex of AD patients. The study identified elevated transcriptome activity and significant changes in lipid metabolism pathways in AD parietal cortex. Notably, non-protein coding isoforms of the *DBI* gene were upregulated in AD. This was one of the earliest studies using RNAseq for AD research, with strengths being the validation of findings by RT-qPCR and sharing of raw data via an online repository. However, the study had several limitations, such as a small sample size, low sequencing depth, potential errors in transcript quantification due to short read lengths (36 nucleotides), lack of a formal AD definition, missing ethnicity data, and lack of females in the study. Furthermore, omission of key information such as the RNA extraction kit used, RNA integrity number, post-mortem interval, and a comprehensive table of differential expression analysis results hinders the study’s rigor and reproducibility.

Guennewig et al.^57^ (published in 2021) analyzed RNAseq data from five AD cases and five controls matched for gender, age, APOE genotype, and RNA integrity number. Patient information included dementia status, clinical dementia rating, and AD pathological diagnosis using the ’ABC’ criteria focused on pathology. All AD cases were Braak stage VI with a high likelihood of AD. Sequencing focused on the precuneus and primary visual cortex. The precuneus showed 559 DEGs, with 462 protein-coding and 97 long non-coding transcripts. The primary visual cortex had 71 DEGs, 44 protein-coding, and 27 long non-coding transcripts, with 40 DEGs shared with the precuneus. Twenty-seven protein-coding DEGs were common to both regions. Eight protein-coding genes were validated by droplet digital PCR, and precuneus DEGs were compared to ROSMAP RNAseq data. Results were also analyzed for overlap with AD risk genes. The study’s strengths include validation using droplet digital PCR and ROSMAP data comparison, sharing raw data in an online repository, and including two brain regions. The main weaknesses were the small sample size, not providing a comprehensive table of differential expression analysis results, and omission of subjects’ ethnicity.

Caldwell et al.^63^ (published in 2022) performed RNAseq on 19 early-onset sporadic AD patients (onset before 60 years) and 20 late-onset AD patients (onset between 70 and 80 years), comparing them with eight aged, non-demented controls. Brain tissue was taken from the primary visual cortex. AD samples were selected based on the absence of alternative diagnoses and APOE status (APOE ε3/3 or ε3/4). The Blessed Information-Memory-Concentration test, the Mini-Mental State Examination, and the Mattis Dementia Rating Scale, along with Braak staging (VI for AD, I or II for controls), were used to classify patients. Hierarchical clustering of RNAseq data identified four transcriptomic clusters, with three clusters mixing early and late-onset cases. One cluster was discarded for quality control. The study found that the number of DEGs compared to controls increased with earlier age of onset and death. Strengths of the study include well-defined AD diagnostic criteria through cognitive and pathological assessments, high sequencing depth, inclusion of both sporadic and late-onset AD cases, and sharing raw data in an online repository. Limitations include a small sample size, no reported attempt at validating results through orthogonal methods or publicly available data, no comprehensive table of differential expression analysis results, and omission of key details such as post-mortem interval and subject ethnicity.

### Studies in multiple brain regions

Out of the 24 literature records included in this systematic review, seven performed bulk RNAseq in multiple brain regions. Twine et al.^46^ (published in 2011) conducted an RNAseq study to compare gene expression and splicing patterns between AD patients and healthy controls, focusing on brain tissues from the temporal lobe, frontal lobe, and whole brain. Their analysis revealed significant differences in genes related to neuronal structure, synapse function, and immune response. Gene ontology analysis highlighted an over-representation of terms linked to synaptic components and neuronal projections, suggesting disruptions in neuronal connectivity and signaling. Notably, they observed changes in the expression levels and promoter usage of the APOE gene, which could provide insight into the mechanisms by which *APOE* isoforms contribute to neurodegeneration. Despite its pioneering use of RNAseq in the year 2011, the study had several limitations. The small sample size, with only one AD sample per brain region, non-age-matched controls with a wide age range, and gender imbalance between groups (females only in the control group) complicated the interpretation of results. Moreover, the study lacked clear criteria for defining AD and omitted key details such as sample ethnicity, RNA extraction methods, RNA integrity numbers, post-mortem intervals, and thresholds for defining differential expression. The use of short-read sequences (36 nucleotides) for transcriptome assembly may have led to inaccuracies in transcript-level abundance estimates, and there were no attempts to validate results with orthogonal techniques.

Miller et al.^45^ (published in 2017) utilized RNAseq to investigate gene expression differences between demented and non-demented individuals from the Adult Changes in Thought study. The study included tissue samples from the temporal lobe, parietal lobe, and hippocampus of 106 participants. The demented group comprised individuals with various forms of dementia, predominantly AD, but also included cases of vascular dementia and mixed dementia. Additionally, half of the subjects had a history of traumatic brain injury.

Surprisingly, the researchers did not identify any DEGs despite employing multiple analysis methods. The study strengths included the comparison of results with previous studies, the availability of raw data in an online repository, the provision of a comprehensive table of differential gene expression results, the inclusion of multiple brain regions, and the reporting of negative findings. The study’s weaknesses included the grouping of diverse dementia types and subjects with traumatic brain injuries into a single "demented" category, which hindered the ability to draw specific conclusions about AD alone.

Lee et al.^54^ (published in 2020) conducted an integrated analysis of H3K9me3 ChIP-seq and RNAseq on cortex tissues of six AD cases and six controls. Diagnoses were confirmed by neuropathologists using NIA Reagan criteria, with AD cases having Braak stage V or VI and controls having Braak stage I or II. The analysis identified 3,367 DEGs, with 1,913 upregulated and 1,454 downregulated in AD. Integrated data revealed 90 genes with inverse relationships between H3K9me3 levels and mRNA expression, including 46 with increased mRNA and decreased H3K9me3 levels and 44 with the opposite pattern. Functional enrichment, network, and gene set enrichment analyses were performed, with results validated by qPCR, western blot, and confocal microscopy. The findings suggest that abnormal heterochromatin remodeling by H3K9me3 leads to the downregulation of synaptic function-related genes, contributing to AD synaptic pathology. Strengths of this study include thorough validation of results using multiple methods, a multi-omics approach combining RNAseq and ChIP-seq, high sequencing depth, and sharing raw data in an online repository. Weaknesses of this study include small sample size, not applying correction for multiple comparisons on p-values, not specifying which cortex tissues were used for RNAseq, not providing demographic information for the subset of samples with RNAseq data, and omitting key details such as RNA integrity number and post-mortem interval.

Li et al.^43^ (published in 2021) compared superior temporal gyrus and inferior frontal gyrus samples from patients with AD and cognitively normal controls. They also investigated differences between patients with mild cognitive impairment and normal controls. The study identifies key genes with altered expression in AD, emphasizing the significance of microglial genes such as *OLR1* and the astrocyte gene *CDK2AP1*. *OLR1* had the strongest association with amyloid plaque burden. *CDK2AP1* had the strongest association with cognitive measures and neurofibrillary tangle burden and also showed an association with amyloid plaque burden.

These findings highlight the critical roles of microglia and astrocytes in the disease’s pathology, potentially offering new targets for therapeutic interventions. Strengths of this study include well-defined AD diagnostic criteria through cognitive and pathological assessments, inclusion of the mild cognitively impaired group, comparison of DEGs with previously published results, and inclusion of multiple brain regions. Weaknesses included not sharing raw data in an online repository, not providing a comprehensive table of differential expression analysis results, and poor age and sex matching between groups.

Marques-Coelho et al.^32^ (published in 2021) does not present original RNAseq data, unlike the other papers included in this systematic review. However, it integrates data from the three largest RNAseq datasets: the Mayo Clinic study^34^ (n=160), the Mount Sinai/JJ Peters VA Medical Center Brain Bank (MSBB)^35^ (n≈165), and the Religious Orders Study and Memory and Aging Project (ROSMAP)^36^ (n=423). To ensure completeness, we included this study to represent these datasets. The study compared RNAseq results from temporal and frontal lobe samples of AD patients and non-demented controls, performing statistical comparisons in each dataset separately before examining similarities and differences in the results between datasets. By combining differential gene expression and isoform switch analyses, the study identified isoform switches in key AD-related genes, such as *APP* and *BIN1*. The differential transcript usage analysis revealed genes undergoing isoform switching without global expression changes. Strengths of this study include a large sample size, multiple brain regions, well-defined AD diagnostic criteria through cognitive and pathological assessments, inclusion of RNA isoform level analysis, single-cell analysis to identify cell-type specific signatures, and the use of western blot to validate some results at the protein level. Raw data is available in an online repository.

Weaknesses included not reporting the ethnicity of subjects—which are mostly Caucasian from the data description publications^34–36^—and the lack of a table providing comprehensive differential gene expression and isoform switching results.

King et al.^55^ (published in 2022) compared mid-life controls and aged controls with AD patients. Aged controls were divided into cognitive resilient and cognitive decline groups for comparison. Total homogenate and synaptoneurosome samples from the middle temporal gyrus and the primary visual cortex of each patient were analyzed. Gene expression involved in synaptic signaling decreased from mid-life to aged and AD cases.

Notably, the cognitive resilient group showed lower expression of synaptic signaling genes compared to the cognitive decline group. Strengths of this study include the use of multiple brain regions, analyzing total brain homogenate and synaptoneurosome separately, well-defined AD diagnostic criteria through cognitive and pathological assessments, a multi-omics approach combining RNAseq and proteomics, and functional validation of results using neuronal cultures. Weaknesses included not sharing raw data in an online repository, assembling the transcriptome with 50 nucleotide short-reads—which could lead to errors in assembly and quantification errors—small sample size, and omitting key methodological details such as sequencing depth.

Santana et al.^51^ (published in 2022) conducted RNAseq in the auditory cortex, hippocampus, and cerebellum of six AD patients and six non-demented controls. Additionally, they performed chromatin immunoprecipitation sequencing (ChIP-Seq) on these same samples using an H3K9 antibody. The researchers identified 22 genes that were both upregulated and hyperacetylated in the auditory cortex. Only one gene, *SCAI*, was found to be simultaneously downregulated and hypoacetylated. Network analysis of these genes revealed involvement in cytoskeletal organization, Rho GTPase-mediated mechanisms, synaptic transmission, and inflammatory processes. Strengths of this study included the use of multiple brain regions, a multi-omics approach combining ChIP-Seq and RNAseq, well-defined AD diagnostic criteria through cognitive and pathological assessments, and high sequencing depth. Limitations of this study included small sample size, imbalanced sex between AD and non-demented control groups, not sharing raw data in an online repository, not providing a comprehensive table of differential expression analysis results, and omitting key details such as subjects’ ethnicity and the software used to generate gene counts matrices.

### Quality Assessment

We assessed the quality of the 24 studies included in this systematic review based on ten categories: "Sample size," "Sex and ethnicity," "AD diagnosis criteria," "Control matching," "Transcript level analysis," "Results validation," "Sequencing depth," "Statistical rigor," "Data availability," and "Reproducibility." Each category was rated on a three-point scale, with one (1) indicating the lowest quality and three (3) indicating the highest quality. The criteria for assigning quality scores are detailed in **Table 2**, and the quality scores for each study are shown in **Table 3**. This quality assessment serves as a proxy for risk of bias assessment in our study. The mean quality score across all studies and categories was 1.8 (0.75). The category with the highest average quality score was "Statistical Rigor," with a mean of 2.4 (0.77), due to most studies correcting p-values for multiple comparisons and/or including covariates in their differential expression analysis models. The category with the lowest average quality score was "Transcript level analysis" with a mean of 1.1 (0.34) due to 21 out of 24 studies not including any transcript level analysis. This omission is significant because statistical analysis at the transcript level has been shown to enhance resolution^65^, despite the limitations of short-read sequencing for measuring expression at the transcript level^66–69^. The study with the highest mean quality score across all categories was Marques-Coelho et al.^32^, with a score of 2.3 (0.48), whereas the study with the lowest mean quality score was Shmookler Reis et al.^46^, with a score of 1.2 (0.68).

**Table 3:** Quality assessment scores for each study. Each study was scored across 10 quality assessment categories, with one (1) being the lowest (worst) score and three (3) being the highest (best) score. Definition of objective criteria for scoring each study can be found in Table 2.

| Study | Sample size | Sex and ethnicity | AD diagnose criteria | Control matching | Transcript level analysis | Results validation | Sequencing depth | Statistical rigor | Data availability | Reproducibility | Average score per study (standard deviation) |
| --- | --- | --- | --- | --- | --- | --- | --- | --- | --- | --- | --- |
| Shmookler Reis et al. | 1 | 1 | 1 | 1 | 1 | 3 | 1 | 1 | 1 | 1 | 1.2 (0.63) |
| Twine et al. | 1 | 1 | 1 | 1 | 2 | 1 | 1 | 2 | 2 | 1 | 1.3 (0.48) |
| Mills et al. | 1 | 1 | 1 | 2 | 2 | 2 | 1 | 2 | 2 | 1 | 1.5 (0.53) |
| Cho et al. | 1 | 2 | 1 | 2 | 1 | 2 | 2 | 2 | 2 | 1 | 1.6 (0.52) |
| Jia et al. | 1 | 2 | 2 | 1 | 1 | 2 | 3 | 2 | 1 | 1 | 1.6 (0.7) |
| Barbash et al. | 1 | 1 | 2 | 2 | 1 | 2 | 3 | 1 | 2 | 1 | 1.6 (0.7) |
| Lee et al. | 1 | 1 | 3 | 1 | 1 | 2 | 3 | 1 | 2 | 1 | 1.6 (0.84) |
| Luo et al. | 2 | 2 | 2 | 1 | 1 | 3 | 1 | 3 | 1 | 1 | 1.7 (0.82) |
| Santana et al. | 1 | 2 | 3 | 1 | 1 | 2 | 3 | 2 | 1 | 1 | 1.7 (0.82) |
| Felsky et al. | 1 | 3 | 2 | 1 | 1 | 2 | 1 | 3 | 2 | 1 | 1.7 (0.82) |
| van Rooij et al. | 1 | 2 | 2 | 2 | 1 | 2 | 2 | 3 | 2 | 1 | 1.8 (0.63) |
| King et al. | 1 | 2 | 3 | 2 | 1 | 3 | 1 | 2 | 2 | 1 | 1.8 (0.79) |
| Humphries et al. | 1 | 1 | 3 | 3 | 1 | 2 | 3 | 2 | 1 | 1 | 1.8 (0.92) |
| Guennewig et al. | 1 | 2 | 2 | 2 | 1 | 2 | 2 | 3 | 2 | 2 | 1.9 (0.57) |
| Li et al. | 2 | 2 | 3 | 1 | 1 | 2 | 2 | 3 | 1 | 2 | 1.9 (0.74) |
| Piras et al. | 1 | 2 | 3 | 1 | 1 | 2 | 2 | 3 | 1 | 3 | 1.9 (0.88) |
| Magistry et al. | 1 | 2 | 3 | 1 | 1 | 3 | 3 | 2 | 2 | 1 | 1.9 (0.88) |
| Fisher et al. | 1 | 2 | 3 | 2 | 1 | 2 | 1 | 3 | 3 | 1 | 1.9 (0.88) |
| Annese et al. | 1 | 1 | 3 | 2 | 1 | 2 | 3 | 2 | 1 | 3 | 1.9 (0.88) |
| Caldwell et al. | 1 | 2 | 3 | 2 | 1 | 1 | 3 | 3 | 2 | 1 | 1.9 (0.88) |
| Das et al. | 1 | 2 | 3 | 2 | 1 | 1 | 2 | 3 | 3 | 2 | 2 (0.82) |
| Panitch et al. | 3 | 2 | 3 | 3 | 1 | 2 | 1 | 3 | 1 | 1 | 2 (0.94) |
| Miller et al. | 2 | 2 | 2 | 2 | 1 | 2 | 2 | 3 | 3 | 2 | 2.1 (0.57) |
| Marques-Coelho et al. | 3 | 2 | 3 | 2 | 2 | 2 | 2 | 3 | 2 | 2 | 2.3 (0.48) |
| Average score per criteria (standard deviation) | 1.3 (0.62) | 1.8 (0.53) | 2.4 (0.77) | 1.7 (0.64) | 1.1 (0.34) | 2 (0.55) | 2 (0.83) | 2.4 (0.71) | 1.8 (0.68) | 1.4 (0.65) | 1.8 (0.75) |

Next, we tested Spearman correlations between quality scores in each category and year of publication (**Supplementary Table 4, Supplementary Figures 1-11**). The quality score correlation for "Sex and Ethnicity" and publication year was the only to reach the Bonferroni corrected p-value < 0.1 threshold (Spearman’s rho = 0.61, p-value = 0.0012), showing a positive correlation with the year of publication. This indicates that more recent publications were more likely to include both sexes in their sample. However, it is important to note that Felsky et al.^61^ was the only publication to include a substantial number of participants from more than one ethnicity. Surprisingly, despite the recent emphasis on improving data availability and reproducibility, there was no correlation between the year of publication and either "Reproducibility" (Spearman’s rho = -0.05, p-value = 0.81) or "Data Availability” (Spearman’s rho = 0.02, p-value = 0.91). Moreover, there was no significant correlation between average quality score across all categories and year of publication (Spearman’s rho = 0.22, p-value = 0.29).

The quality assessment scale presented here can help guide the design of future transcriptomic studies in AD using technologies such as single-cell RNAseq, long-read RNAseq, spatial transcriptomics, and epitranscriptomics. Although some criteria—like sequencing depth in single-cell RNAseq studies—may require updates, most of the scoring system should remain applicable across various technologies.

### Meta-analysis overview

We conducted a meta-analysis using three datasets examined by Marques-Coelho and colleagues^32^: Mayo Clinic temporal lobe^34^, MSBB frontal and temporal lobes^35^, and ROSMAP dorsolateral prefrontal cortex^36^.

These datasets were selected based on three key factors: first, each contains over 150 samples, which is significantly larger than the next largest study that provided a table suitable for meta-analysis, which included 106 samples. Second, the study analyzing these datasets achieved the highest quality assessment score, thereby minimizing the risk of bias. Third, among the remaining 23 studies, only five provided differential expression tables suitable for meta-analysis; however, these were excluded due to either small sample sizes (n < 50) or substantial methodological concerns, such as loosely defined AD status in the study with 106 samples. We split these datasets into two separate meta-analyses. The first was done on the temporal lobe, using the MSBB Brodmann area 22 dataset (n = 159) and the Mayo temporal lobe data from Brodmann areas 20/21/22/41/42 (n = 160). The second meta-analysis was done on the frontal lobe, utilizing the MSBB Brodmann area 10 dataset (n = 176) and ROSMAP Brodmann areas 9/46 (n = 423). We excluded two MSBB datasets—Brodmann area 44 (frontal lobe) and Brodmann area 36 (temporal lobe)—due to overlapping samples with the included datasets, which would violate the data independence assumption for our meta-analysis. MSBB Brodmann area 22 was chosen for the temporal lobe analysis over Brodmann area 36 due to its slightly larger sample size and greater anatomical and functional similarity to the Mayo temporal lobe dataset regions. For the frontal lobe analysis, MSBB Brodmann area 10 was selected over Brodmann area 44 for its slightly larger sample size and closer anatomical and functional resemblance to the ROSMAP dataset’s Brodmann areas.

### Meta-analysis in temporal lobe

We performed a differential gene expression meta-analysis on two temporal lobe datasets analyzed by Marques-Coelho and colleagues^32^. The datasets included MSBB Brodmann area 22 (n = 159) and Mayo temporal lobe data (n = 160), totaling 319 samples. Genes were considered differentially expressed between AD and non-demented controls if the Bonferroni adjusted p-value was < 0.1 (equivalent to unadjusted p-value < 3.39 × 10^−6^). Using this threshold, we identified 418 upregulated genes in AD and 153 downregulated genes, resulting in 571 total DEGs (**Figure 2; Supplementary Tables 5-7).** Out of these 571 DEGs, 41 were uniquely identified in our meta-analysis, distinguishing them from DEGs in eight other temporal lobe studies reviewed; 10 DEGs were shared with five of these studies (**Table 4; Supplementary Table 8**). The I^2^ metric for heterogeneity was moderate, averaging 25.22% (SD = 34.32%) across the temporal lobe meta-analysis (**Supplementary Table 9**).

**Figure 2:**
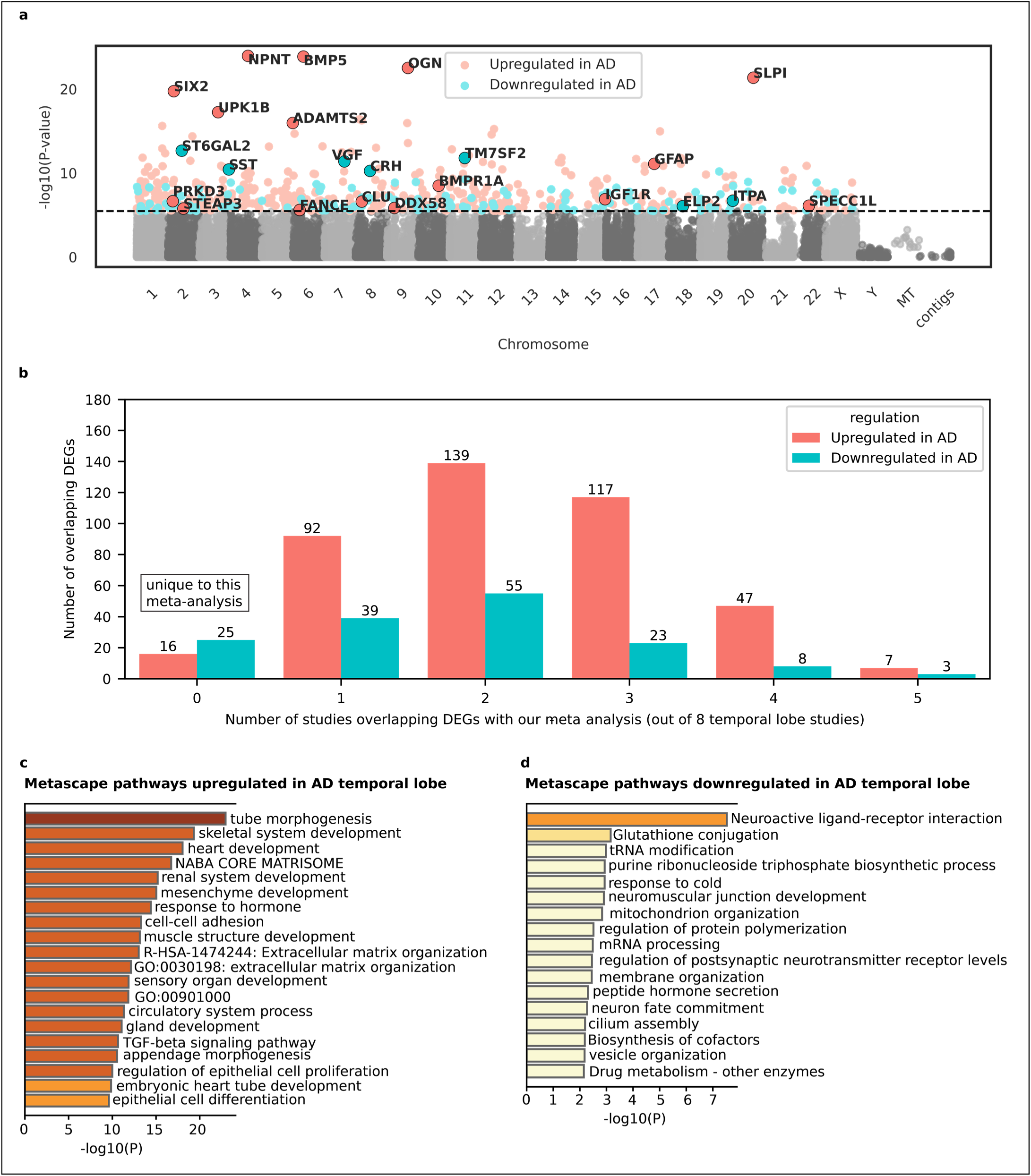
Meta-analysis for AD temporal lobe differential gene expression. **a,** Differential gene expression results for the AD temporal lobe meta-analysis (n = 319; 180 AD and 139 controls). The Manhattan plot shows - log10(p-value) on the y-axis and chromosome coordinates on the x-axis. The dotted line represents the Bonferroni corrected p-value threshold of 0.1 (equivalent to an unadjusted p-value of 3.39 × 10^−6^). Genes were considered significantly differentially expressed between AD subjects and controls if the Bonferroni corrected p-value was less than to 0.1. Statistics were derived from the Inverse-Variance Weighted Fixed-Effect Model from METAL^37^. Red dots indicate genes upregulated in AD, while blue dots indicate genes downregulated in AD. Selected differentially expressed genes are highlighted with their gene symbols for interest. **b,** Number of temporal lobe meta-analysis DEGs overlapping with other studies. The y-axis represents the number of overlapping DEGs, while the x-axis indicates how many of the eight temporal lobe studies each DEG overlapped with. A value of "0" on the x-axis corresponds to DEGs that are unique to this meta-analysis. More detailed information about DEG overlap for temporal lobe can be found in **Supplementary Table 8**. **c,** Metascape pathway convergence analysis for genes upregulated in the AD temporal lobe. Pathways are listed on the y-axis, with -log10(p-value) on the x-axis. **d,** Metascape pathway convergence analysis for genes downregulated in the AD temporal lobe. Pathways are listed on the y-axis, with -log10(p-value) on the x-axis.

**Table 4:**
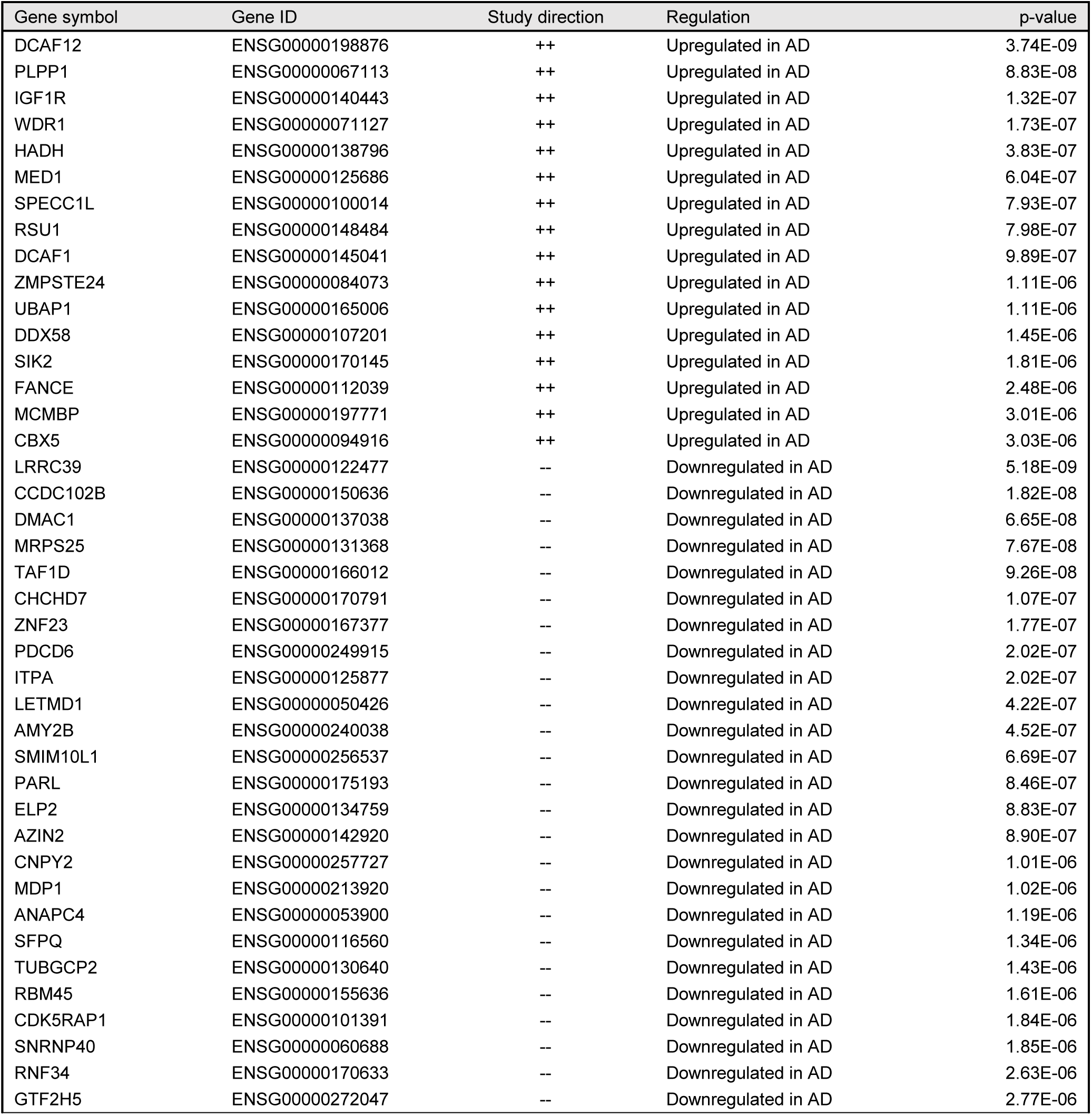
Temporal lobe meta-analysis DEGs unique to our study. List of differentially expressed genes in temporal lobe meta-analysis that were unique to our study when compared to 8 other temporal lobe studies included in this systematic review. P-value derived from the Inverse-Variance Weighted Fixed-Effect Model from METAL (n=319).

| Gene symbol | Gene ID | Study direction | Regulation | p-value |
| --- | --- | --- | --- | --- |
| DCAF12 | ENSG00000198876 | ++ | Upregulated in AD | 3.74E-09 |
| PLPP1 | ENSG00000067113 | ++ | Upregulated in AD | 8.83E-08 |
| IGF1R | ENSG00000140443 | ++ | Upregulated in AD | 1.32E-07 |
| WDR1 | ENSG00000071127 | ++ | Upregulated in AD | 1.73E-07 |
| HADH | ENSG00000138796 | ++ | Upregulated in AD | 3.83E-07 |
| MED1 | ENSG00000125686 | ++ | Upregulated in AD | 6.04E-07 |
| SPECC1L | ENSG00000100014 | ++ | Upregulated in AD | 7.93E-07 |
| RSU1 | ENSG00000148484 | ++ | Upregulated in AD | 7.98E-07 |
| DCAF1 | ENSG00000145041 | ++ | Upregulated in AD | 9.89E-07 |
| ZMPSTE24 | ENSG00000084073 | ++ | Upregulated in AD | 1.11E-06 |
| UBAP1 | ENSG00000165006 | ++ | Upregulated in AD | 1.11E-06 |
| DDX58 | ENSG00000107201 | ++ | Upregulated in AD | 1.45E-06 |
| SIK2 | ENSG00000170145 | ++ | Upregulated in AD | 1.81E-06 |
| FANCE | ENSG00000112039 | ++ | Upregulated in AD | 2.48E-06 |
| MCMBP | ENSG00000197771 | ++ | Upregulated in AD | 3.01E-06 |
| CBX5 | ENSG00000094916 | ++ | Upregulated in AD | 3.03E-06 |
| LRRC39 | ENSG00000122477 | -- | Downregulated in AD | 5.18E-09 |
| CCDC102B | ENSG00000150636 | -- | Downregulated in AD | 1.82E-08 |
| DMAC1 | ENSG00000137038 | -- | Downregulated in AD | 6.65E-08 |
| MRPS25 | ENSG00000131368 | -- | Downregulated in AD | 7.67E-08 |
| TAF1D | ENSG00000166012 | -- | Downregulated in AD | 9.26E-08 |
| CHCHD7 | ENSG00000170791 | -- | Downregulated in AD | 1.07E-07 |
| ZNF23 | ENSG00000167377 | -- | Downregulated in AD | 1.77E-07 |
| PDCD6 | ENSG00000249915 | -- | Downregulated in AD | 2.02E-07 |
| ITPA | ENSG00000125877 | -- | Downregulated in AD | 2.02E-07 |
| LETMD1 | ENSG00000050426 | -- | Downregulated in AD | 4.22E-07 |
| AMY2B | ENSG00000240038 | -- | Downregulated in AD | 4.52E-07 |
| SMIM10L1 | ENSG00000256537 | -- | Downregulated in AD | 6.69E-07 |
| PARL | ENSG00000175193 | -- | Downregulated in AD | 8.46E-07 |
| ELP2 | ENSG00000134759 | -- | Downregulated in AD | 8.83E-07 |
| AZIN2 | ENSG00000142920 | -- | Downregulated in AD | 8.90E-07 |
| CNPY2 | ENSG00000257727 | -- | Downregulated in AD | 1.01E-06 |
| MDP1 | ENSG00000213920 | -- | Downregulated in AD | 1.02E-06 |
| ANAPC4 | ENSG00000053900 | -- | Downregulated in AD | 1.19E-06 |
| SFPQ | ENSG00000116560 | -- | Downregulated in AD | 1.34E-06 |
| TUBGCP2 | ENSG00000130640 | -- | Downregulated in AD | 1.43E-06 |
| RBM45 | ENSG00000155636 | -- | Downregulated in AD | 1.61E-06 |
| CDK5RAP1 | ENSG00000101391 | -- | Downregulated in AD | 1.84E-06 |
| SNRNP40 | ENSG00000060688 | -- | Downregulated in AD | 1.85E-06 |
| RNF34 | ENSG00000170633 | -- | Downregulated in AD | 2.63E-06 |
| GTF2H5 | ENSG00000272047 | -- | Downregulated in AD | 2.77E-06 |

Two genes previously implicated in AD through genome-wide association studies^9^ were upregulated in the temporal lobe of AD samples: *PRKD3* (Serine/Threonine-Protein Kinase D3; involved in vesicle transport^70^), and *CLU* (involved in beta-amyloid clearance^71,72^). A notable downregulated gene was *SST* (encodes the peptide hormone somatostatin, involved in modulating glutamatergic signals in the central nervous system^73^). *HADH*, a key enzyme involved in converting medium- and short-chain fatty acids into ketones—a crucial alternative energy source for the brain during glucose deprivation—was found to be upregulated in AD in our meta-analysis. This enzyme, also known for binding amyloid-beta peptides^74^, was uniquely identified in our study compared to eight other temporal lobe studies. *PARL*, a serine protease previously linked to Parkinson’s disease^75^, was downregulated and also uniquely identified in our meta-analysis. *GAD1*, which encodes the enzyme responsible for producing the gamma-aminobutyric acid (GABA) neurotransmitter^76,77^, was downregulated not only in our meta-analysis but also in five other temporal lobe studies. Similarly, *STEAP3*, a gene involved in the ferroptosis pathway^78,79^, was upregulated in both our meta-analysis and in five other temporal lobe studies.

The Metascape analysis for upregulated genes revealed pathways converging around developmental processes (gland development, mesenchyme development, sensory organ development, muscle structure development, heart development, renal system development, and embryonic heart tube development), suggesting aberrant developmental pathway reactivation in the AD brain. Metascape analysis for downregulated genes in AD temporal lobe produced less robust results compared to upregulated genes. A couple pathways related to neuronal function (Neuroactive ligand-receptor interaction and regulation of postsynaptic membrane neurotransmitter receptor levels) were downregulated in AD, indicating neuronal death and/or dysfunction as possible drivers of differential gene expression.

### Meta-analysis in frontal lobe

We performed differential gene expression meta-analysis on two frontal lobe datasets analyzed by Marques- Coelho and colleagues^32^. The datasets included MSBB Brodmann area 10 (n = 176) and ROSMAP Brodmann area 9/46 (n = 423), totaling 599 samples. Genes were considered differentially expressed between AD and non-demented controls if the Bonferroni adjusted p-value was < 0.1 (equivalent to unadjusted p-value 3.19 × 10^−6^). Using this threshold, we found 154 genes upregulated in AD and 35 downregulated genes, for a total of 189 DEGs (**Figure 3; Supplementary Table 10-12**). Out of these 189 DEGs, 39 were uniquely identified in our meta-analysis, distinguishing them from DEGs in three other frontal lobe studies reviewed; 10 DEGs were shared with three of these studies (**Table 5; Supplementary Table 13**). The I^2^ metric for heterogeneity was low, averaging 9.79% (SD = 21.84%) across genes in the frontal lobe meta-analysis (**Supplementary Table 3**).

**Figure 3:**
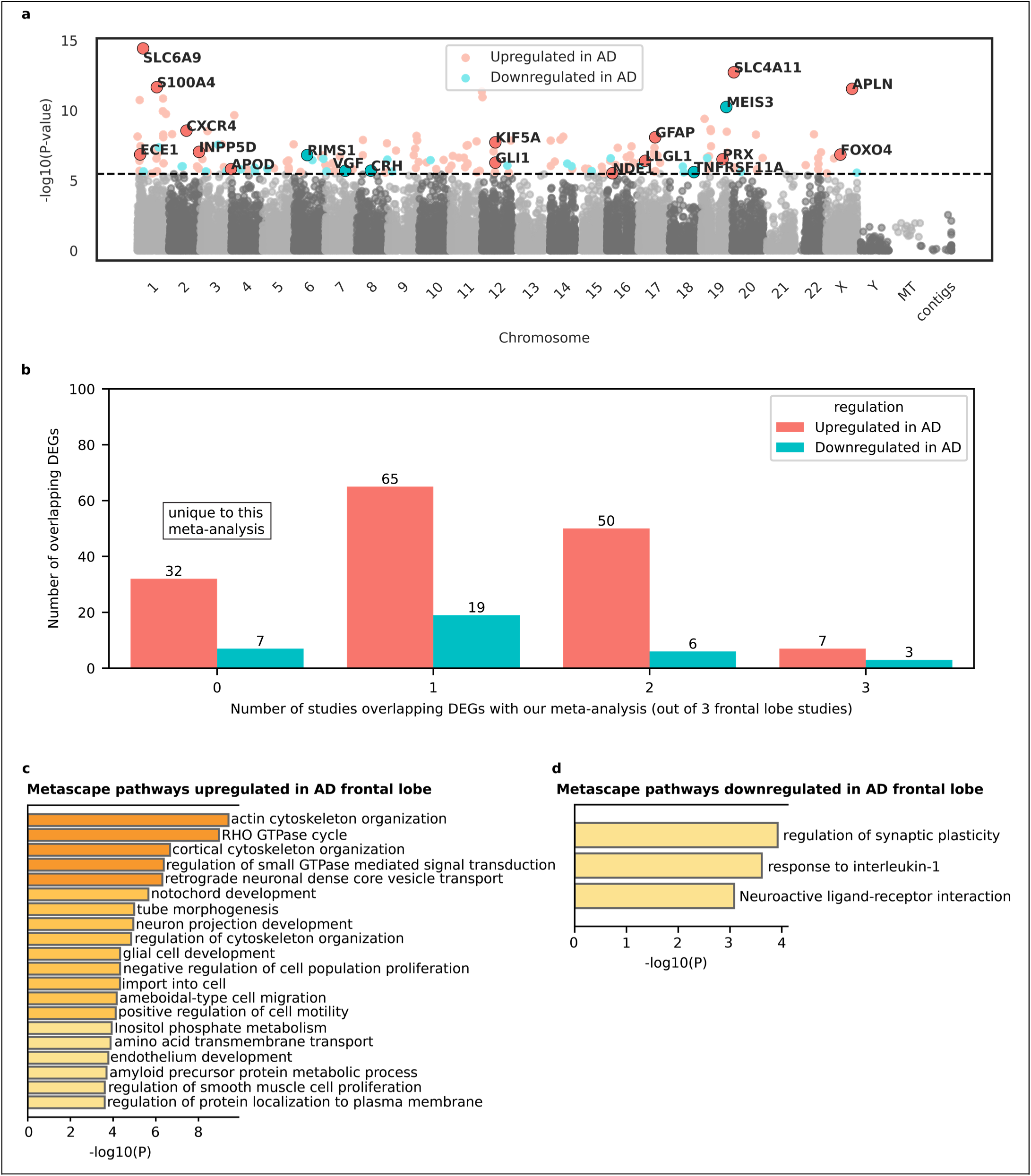
Meta-analysis for AD frontal lobe differential gene expression. **a,** Differential gene expression results for the AD frontal lobe meta-analysis (n = 599; 327 AD and 272 controls). The Manhattan plot shows - log10(p-value) on the y-axis and chromosome coordinates on the x-axis. The dotted line represents the Bonferroni corrected p-value threshold of 0.1 (equivalent to a unadjusted p-value of 3.19 × 10^−6^.). Genes were considered significantly differentially expressed between AD subjects and controls if the Bonferroni corrected p-value was less than 0.1. Statistics were derived from the Inverse-Variance Weighted Fixed-Effect Model from METAL^37^. Red dots indicate genes upregulated in AD, while blue dots indicate genes downregulated in AD. Selected differentially expressed genes are highlighted with their gene symbols. **b,** Number of frontal lobe meta-analysis DEGs overlapping with other studies. The y-axis represents the number of overlapping DEGs, while the x-axis indicates how many of the three frontal lobe studies each DEG overlapped with. A value of "0" on the x-axis corresponds to DEGs that are unique to this meta-analysis. **c,** Metascape pathway convergence analysis for genes upregulated in the AD frontal lobe. Pathways are listed on the y-axis, with -log10(p-value) on the x-axis. **d,** Metascape pathway convergence analysis for genes downregulated in the AD frontal lobe. Pathways are listed on the y-axis, with -log10(p-value) on the x-axis.

**Table 5:**
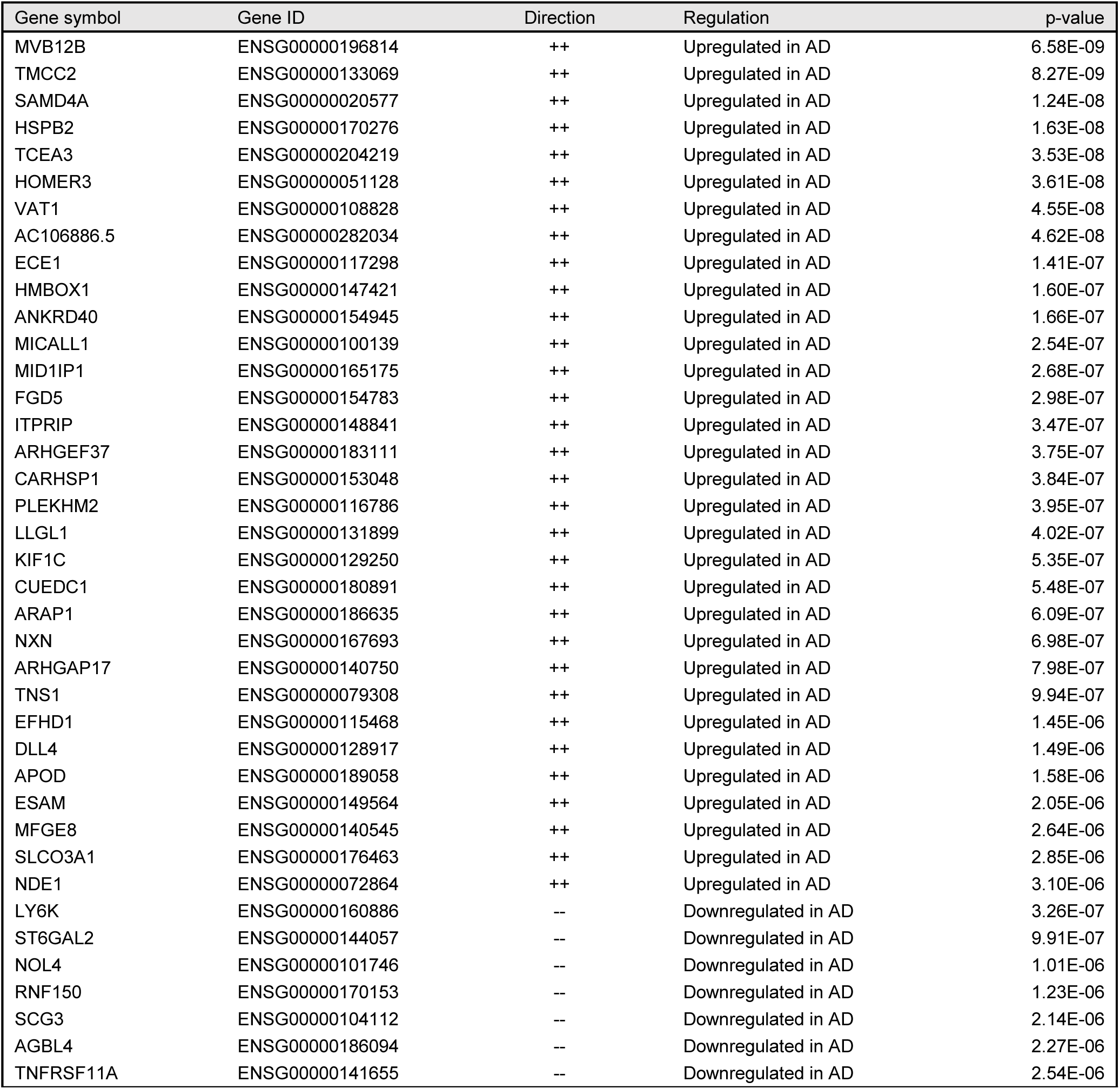
Frontal lobe meta-analysis DEGs unique to our study. List of differentially expressed genes in frontal lobe meta-analysis that were unique to our study when compared to 3 other frontal lobe studies included in this systematic review. P-value derived from the Inverse-Variance Weighted Fixed-Effect Model from METAL (n=599).

*FOXO4*, a transcription factor involved in cellular senescence, oxidative stress, and metabolism^80,81^, is upregulated in the AD frontal cortex but not in the temporal cortex. *FOXO4* has been shown to inhibit *HIF1A*^82,83^, a transcription factor linked to AD-related microglia phenotypic changes in both mouse and human cell-line models^84,85^. Other upregulated genes in AD frontal lobe included *INPP5D* (AD risk gene^9^; regulates inflammasome activation and autophagy in human microglia^86^), *PRX* (involved in peripheral myelin upkeep^87^), and *KIF5A* (a causal gene for familial amyotrophic lateral sclerosis^88,89^). A notable downregulated gene in AD frontal cortex, but not in the temporal cortex, was *RIMS1* which encodes three RNA isoforms integral to modulating synaptic vesicle fusion and presynaptic plasticity^90^. *CXCR4*, a chemokine receptor implicated in microglial responses to neurodegenerative diseases^91^ was upregulated in AD in our meta-analysis as well as three other frontal lobe studies.

The Metascape pathway analysis results for the frontal lobe were less robust than those for the temporal lobe. They revealed upregulated pathways centered on nervous system development (glial cell development, neuron projection development, and notochord development) and cytoskeletal and structural changes (Regulation of cytoskeleton organization, actin cytoskeleton organization, cortical cytoskeleton organization, tube morphogenesis, and RHO GTPase cycle). An important pathway for AD, the amyloid precursor protein metabolic process, was upregulated in the frontal lobe but not in the temporal lobe. These findings highlight aberrant developmental signatures upregulated in the adult AD brain along with increases in cytoskeletal structural changes. Metascape analysis for downregulated genes in AD frontal lobe produced less robust results compared to upregulated genes. Two pathways related to neuronal function—regulation of synaptic plasticity and neuroactive ligand-receptor interaction—were downregulated in AD, again suggesting that neuronal death and/or dysfunction may contribute to the observed differential gene expression in the frontal lobe.

### Temporal and frontal lobe convergence

There was some overlap between differentially expressed genes in the temporal and frontal lobe meta-analyses (**Supplementary Table 14).** Of the 189 differentially expressed genes in the frontal lobe, 44 (23.3%) overlapped with those in the temporal lobe. Specifically, 38 out of 154 (24.7%) upregulated genes and 6 out of 35 (17.1%) downregulated genes were shared. The most notable shared upregulated gene was *GFAP* (glial fibrillary acidic protein; involved in astrocyte reactivity^92,93^), a promising biomarker for AD, with increased expression observed in corticospinal fluid and blood of AD patients^94–98^. *VGF* was a notable downregulated gene, currently being studied as a potential biomarker for AD, with reduced expression observed in AD patient corticospinal fluid samples^99,100^. Additionally, *CRH* (corticotropin-releasing hormone) involved in the allostatic stress response and learning^101^, is also downregulated in temporal and frontal lobe.

Among the upregulated Metascape pathways, "tube morphogenesis" was common to both the frontal and temporal lobes, suggesting a consistent role in AD pathology. This pathway, crucial for tissue structural development, indicates potential alterations in tissue organization and integrity or the presence of aberrant developmental signatures in adult AD brains. Among the downregulated pathways, "Neuroactive ligand-receptor interaction" was observed in both lobes. This pathway is essential for neuronal communication and its downregulation implies disruption in synaptic activity and neurotransmitter interactions, possibly caused by neuronal dysfunction or death in AD.

## Discussion

This systematic review presents a comprehensive catalog of bulk RNAseq studies that use original data to compare human AD brains with non-demented controls. We briefly describe the main findings from the 24 included studies, highlighting their strengths and limitations. This catalog serves as a valuable resource for AD researchers seeking to identify gaps in the literature or build on previous ideas and study designs. Additionally, we developed a quality assessment scale based on 10 objective criteria and applied it to these 24 studies, highlighting key considerations for designing the next generation of transcriptomic studies in AD. Finally, we expanded upon the RNAseq analysis conducted by Marques-Coelho et al.^32^ by performing a meta-analysis on the frontal and temporal lobes separately, then comparing the results. Our approach offers insights into the transcriptomic changes occurring in the AD brain.

Our quality assessment tool for AD transcriptomics studies comprises 10 categories with objective criteria, each scored from one to three, where one represents the lowest quality and three the highest. Detailed descriptions of these criteria are provided in **Table 2**. A key area for improvement identified by our assessment is data availability—specifically, the need to deposit raw data in online repositories and provide comprehensive tables with differential expression analysis results. Addressing these issues would enhance the available datasets for the broader AD research community and facilitate cross-study comparisons, potentially uncovering critical differences and common themes in AD transcriptomic research. While initiatives like the AD Knowledge Portal^33^ have improved data accessibility, significant gaps remain. Additionally, we emphasize the lack of diversity in the populations studied. While women’s representation has improved in recent years (**Supplementary Table 4, Supplementary Figure 2**), only one study included a substantial number of subjects from multiple ancestry backgrounds, with all others either failing to report on participant ethnicity or focusing exclusively on subjects of European or Chinese ancestry. Researchers must prioritize recruiting participants from diverse ancestries into brain banks and AD research cohorts. This is essential for advancing our understanding of AD across different ancestry backgrounds, particularly given the disease’s strong genetic influences. For instance, a recent genome-wide association study found that APOE genotypes^102^—the largest single risk factor for late-onset AD—exhibit significantly different effect sizes across ancestries such as East Asian, non-Hispanic white, non-Hispanic Black, and Hispanic. This underscores the urgent need to study diverse ancestries to fully understand AD mechanisms.

Our quality assessment highlighted the necessity for more thorough methodological reporting to improve research reproducibility and the need for larger sample sizes with greater statistical power, as 19 of the 24 studies had sample sizes smaller than 50. Another area for improvement is the inclusion of transcript-level differential expression or differential usage analysis, which was performed in only three out of the 24 studies. This shortfall is likely due to the limitations of short-read sequencing in accurately quantifying and assembling RNA isoforms, given the high homology between isoforms of the same gene. This homology increases the uncertainty in quantification and assembly when using short reads^66–69^, leading researchers to favor gene-level analysis that collapses all RNA isoforms into a single measurement—a major oversimplification of the underlying biology. However, despite these limitations, work by Love et al.^65^ demonstrates that transcript-level analysis enhances resolution for short-read RNAseq differential expression analysis. Therefore, while it is advisable to proceed with caution and validate key RNA isoform results using orthogonal methods, short-read RNAseq studies would benefit from incorporating differential RNA isoform usage and expression analysis— especially those utilizing 100-nucleotide or longer paired-end short-read sequencing approaches.

Our meta-analysis revealed intriguing patterns of differential gene expression between AD cases and controls. We observed some overlap between differentially expressed genes in the frontal and temporal lobes. Notably, two genes—*GFAP* and *VGF*—both of which have shown promise as peripheral biomarkers for AD, were significantly differentially expressed in both brain regions. Importantly, these genes exhibited the same differential expression pattern as their peripheral biomarkers, with *GFAP* upregulated in AD^94–98^ and *VGF* downregulated in AD^99,100^. This finding underscores the potential of RNAseq in the brain to identify potential peripheral biomarkers for AD. Additionally, *FOXO4* was upregulated in the AD frontal cortex but not in the temporal cortex. *FOXO4* has been shown to inhibit *HIF1A*^82,83^, a transcription factor linked to microglial phenotypic changes in AD models using cell lines and mice^84,85^. This differential expression pattern may suggest an early compensatory mechanism to balance microglial responses to beta-amyloid, which could be lost in the more advanced pathology typically seen in the temporal lobe. Our meta-analysis also revealed that upregulated genes in the temporal and frontal lobes both present Metascape terms related to developmental signatures, highlighting aberrant developmental activation in the adult AD brain. Together, these findings demonstrate that RNAseq is a powerful tool that can uncover putative disease mechanisms. However, follow-up experiments are essential to validate promising targets identified through RNAseq. The associative nature of human RNA-seq studies makes it difficult to distinguish gene expression changes that are a result of AD from those that contribute to its development. While both findings hold value—the former for biomarker identification and the latter for therapeutic advances—a deeper understanding of these causal relationships is crucial for meaningful downstream applications.

Recently, new technologies beyond bulk short-read RNAseq have shown great promise in advancing our understanding of transcriptomic changes in AD. Long-read RNAseq by PacBio and Oxford Nanopore Technologies improves the quantification and assembly for RNA isoforms^66–69^, enabling more accurate differential isoform expression studies. For instance, a pilot study by Heberle et al.^66^ used long-read RNAseq and found 99 RNA isoforms differentially expressed between AD subjects and controls, even when the overall gene was not differentially expressed. However, larger studies are needed to better understand associations between RNA isoform expression and AD. In addition, single-cell and single-nucleus RNAseq have identified important cell-specific RNA programs in AD, such as disease-associated microglia^103^ and other cell-type specific changes^104–109^—opening new avenues for therapeutic interventions. The next step will be to combine long-read RNAseq with single-cell techniques to explore cell-type specific RNA isoform changes in AD. However, this is currently limited by the prohibitive costs of combining both techniques and challenges in performing true single-cell preparations for human brain tissue. Most studies in human post-mortem brain tissue rely on single-nuclei rather than single-cell preparations, which are not so effective for RNA isoform studies since most nuclear mRNAs are unspliced pre-mRNAs. Achieving the full benefits of long-read sequencing requires true single-cell long-read sequencing in post-mortem brain tissue, which is difficult due to the need for fresh tissue and the brain cell fragility during single-cell preparation.

Spatial transcriptomics platforms such as Visium by 10X genomics are promising in AD research. By creating precise gene expression maps in the human brain, they could help us understand why certain brain regions are more vulnerable to AD pathology. This approach has already started to yield results, with a study identifying layer-specific DEGs in the human middle temporal gyrus between AD subjects and controls^110^.

Spatial transcriptomics cell-type deconvolution using single-cell references may provide further insights into the cellular composition of vulnerable regions such as the hippocampus and entorhinal cortex. Another major innovation comes from direct RNAseq by Oxford Nanopore Technologies. By sequencing RNA molecules in their native form and bypassing the need for conversion into cDNA, this technology can detect modifications such as N6-Methyladenosine, which plays an essential role in mRNA maturation, translation, and decay^111^. By retaining these epitranscriptomic modifications, direct RNAseq provides a more comprehensive view of the transcriptome, enabling RNA modification detection that was previously inaccessible with cDNA-based methods. This approach holds great promise for uncovering insights into the molecular mechanisms underlying AD. However, further development is needed to reliably quantify a broader range of RNA modifications before the technology can fully reach its potential^112–116^.

This systematic review and meta-analysis has limitations, however. Due to the large number of studies analyzing the bulk short-read RNAseq data from the AD Knowledge Portal—specifically the Mayo Clinic study^34^, MSBB^35^, and ROSMAP^36^—we excluded studies that did not report original RNAseq data. The exception was the Marques-Coelho et al.^32^ study, which we considered the best representation of these three datasets in a single study, examining both gene-level and transcript-level differential expression/usage.

However, it is possible that we missed key findings from studies that employed different approaches to analyzing these same data. Our meta-analysis was limited by the scarcity of articles with comprehensive differential expression analysis tables and the small sample sizes or loose definitions of AD in the five studies that included such tables. Consequently, we proceeded with the meta-analysis focusing solely on the large datasets previously analyzed by Marques-Coelho and colleagues^32^. Despite the larger sample size, our results would have been more robust if more studies with large sample sizes had provided comprehensive differential expression analysis tables. Additionally, the MSBB datasets included in our meta-analysis were not independent, as they originated from different brain regions of overlapping subjects. To mitigate this issue, we only included one MSBB dataset in the meta-analysis for each brain region. However, it is possible that the overlap in DEGs between the frontal and temporal lobes is partly due to the dependence between MSBB datasets. Our meta-analysis approach does not report effect sizes, such as log2FoldChange. Consequently, we did not filter DEGs based on effect sizes, which may result in the inclusion of DEGs unique to our analysis, as they might have been excluded from other studies that applied effect size thresholds. In addition, without an effect size filter, some DEGs in our results may exhibit small effect sizes. Lastly, the use of Bonferroni correction in a meta-analysis that used Benjamini-Hochberg corrected p-values as input may have led to false negatives due to the stringent significance thresholds.

The information synthesis presented in this systematic review and meta-analysis can assist researchers in identifying gaps in the literature, allowing them to build on previous ideas and refine study designs for future research. The design and execution of next-generation transcriptomic studies in AD—including single-cell, long-read, direct RNAseq, and spatial transcriptomics—can benefit substantially from the quality assessment criteria introduced in this review. By applying these criteria to both experiment design and reporting, researchers using these technologies will be able to produce more robust results characterized by greater rigor, reproducibility, and generalizability. While certain categories, such as “sequencing depth,” may require adaptation for specific contexts like single-cell experiments, many other criteria—such as “AD diagnostic criteria,” “reproducibility,” “data availability,” “results validation,” and considerations of “sex and ethnicity”—are broadly applicable across various transcriptomic study designs. Overall, integrating these criteria into transcriptomic studies will not only enhance the quality and reliability of the findings but also improve the likelihood that the resulting data can eventually be effectively translated into clinical applications. Lastly, the meta-analysis performed here provides a different look into AD transcriptomic changes, highlighting possible biomarkers and therapeutic targets.

## Data availability

Additional files such as meta-analysis input files, meta-analysis output files, and full Metascape pathway analysis output were deposited in an open Zenodo repository: https://zenodo.org/records/13754371.

## Code availability

All code used in the manuscript and the instructions to retrieve the Singularity containers used to execute the code are publicly available at https://github.com/UK-SBCoA-EbbertLab/AD_bulk_RNAseq_review.

## Supporting information

Supplementary File 1 - PRISMA Checklist

Supplementary File 2 - Prisma Abstract Checklist

## Acknowledgments

This work was supported by the National Institutes of Health [R35GM138636, R01AG068331] to M.T.W.E., [RF1AG082339, P30AG072946] to M.T.W.E. and D.W.F., and [R01AG082730] to D.W.F. Additional support was provided by the BrightFocus Foundation [A2020161S to M.T.W.E.], Alzheimer’s Association [2019-AARG44082 to M.T.W.E.], PhRMA Foundation [RSGTMT17 to M.T.W.E.]; Ed and Ethel Moore Alzheimer’s Disease Research Program of Florida Department of Health [8AZ10 and 9AZ08 to M.T.W.E.]; and the Muscular Dystrophy Association (M.T.W.E.). We appreciate the contributions of the Sanders-Brown Center on Aging at the University of Kentucky. We are deeply grateful to the research participants and their families who make this research possible. We would like to thank the University of Kentucky Center for Computational Sciences and Information Technology Services Research Computing for their support and use of the Morgan Compute Cluster and associated research computing resources. We would like to thank Singularity Sylabs for providing support and extra cloud storage for our software containers.

## Author Contributions Statement

B.A.H., K.L.F., T.W.V., and M.T.W.E. developed and designed the study and wrote the paper. B.A.H. and L.L.L. performed the database searches and eliminated record duplicates. B.A.H., K.L.F., L.L.L., S.R., G.D. and R.S. performed the title and abstract screening. B.A.H. and K.L.F. performed the full-text screening, data extraction, and quality assessment. B.A.H. performed all analyses. D.W.F was instrumental in developing the meta-analysis plan and made contributions to the manuscript writing.

## References

1. 2023 Alzheimer’s disease facts and figures - 2023 - Alzheimer’s & Dementia - Wiley Online Library. https://alz-journals.onlinelibrary.wiley.com/doi/10.1002/alz.13016.

2. GBD 2019 Dementia Forecasting Collaborators. Estimation of the global prevalence of dementia in 2019 and forecasted prevalence in 2050: an analysis for the Global Burden of Disease Study 2019. Lancet Public Health **7**, e105– e125 (2022).

3. Dementia. https://www.who.int/news-room/fact-sheets/detail/dementia.

4. Rajan, K. B. et al. Population estimate of people with clinical Alzheimer’s disease and mild cognitive impairment in the United States (2020-2060). Alzheimers Dement. J. Alzheimers Assoc. 17, 1966–1975 (2021).

5. Weller, J. & Budson, A. Current understanding of Alzheimer’s disease diagnosis and treatment. F1000Research **7**, F1000 Faculty Rev-1161 (2018).

6. Tatulian, S. A. Challenges and hopes for Alzheimer’s disease. Drug Discov. Today 27, 1027–1043 (2022).

7. Bergem, A. L., Engedal, K. & Kringlen, E. The role of heredity in late-onset Alzheimer disease and vascular dementia. A twin study. Arch. Gen. Psychiatry 54, 264–270 (1997).

8. Gatz, M. et al. Role of genes and environments for explaining Alzheimer disease. Arch. Gen. Psychiatry 63, 168– 174 (2006).

9. Bellenguez, C. et al. New insights into the genetic etiology of Alzheimer’s disease and related dementias. Nat. Genet. 54, 412–436 (2022).

10. Shade, L. M. P. et al. GWAS of multiple neuropathology endophenotypes identifies new risk loci and provides insights into the genetic risk of dementia. Nat. Genet. 1–15 (2024) doi:10.1038/s41588-024-01939-9.

11. Golde, T. E., Estus, S., Usiak, M., Younkin, L. H. & Younkin, S. G. Expression of beta amyloid protein precursor mRNAs: recognition of a novel alternatively spliced form and quantitation in Alzheimer’s disease using PCR. Neuron 4, 253–267 (1990).

12. Neve, R. L., Rogers, J. & Higgins, G. A. The Alzheimer amyloid precursor-related transcript lacking the beta/A4 sequence is specifically increased in Alzheimer’s disease brain. Neuron 5, 329–338 (1990).

13. Johnson, S. A., McNeill, T., Cordell, B. & Finch, C. E. Relation of neuronal APP-751/APP-695 mRNA ratio and neuritic plaque density in Alzheimer’s disease. Science 248, 854–857 (1990).

14. Golde, T. E., Eckman, C. B. & Younkin, S. G. Biochemical detection of Aβ isoforms: implications for pathogenesis, diagnosis, and treatment of Alzheimer’s disease. Biochim. Biophys. Acta BBA - Mol. Basis Dis. 1502, 172–187 (2000).

15. Goedert, M., Wischik, C. M., Crowther, R. A., Walker, J. E. & Klug, A. Cloning and sequencing of the cDNA encoding a core protein of the paired helical filament of Alzheimer disease: identification as the microtubule-associated protein tau. Proc. Natl. Acad. Sci. 85, 4051–4055 (1988).

16. Goedert, M., Spillantini, M. G., Potier, M. C., Ulrich, J. & Crowther, R. A. Cloning and sequencing of the cDNA encoding an isoform of microtubule-associated protein tau containing four tandem repeats: differential expression of tau protein mRNAs in human brain. EMBO J. 8, 393–399 (1989).

17. Andreadis, A., Brown, W. M. & Kosik, K. S. Structure and novel exons of the human tau gene. Biochemistry 31, 10626–10633 (1992).

18. Goedert, M., Crowther, R. A. & Garner, C. C. Molecular characterization of microtubule-associated proteins tau and map2. Trends Neurosci. 14, 193–199 (1991).

19. Conrad, C. et al. Single molecule profiling of tau gene expression in Alzheimer’s disease. J. Neurochem. 103, 1228–1236 (2007).

20. Ho, L. et al. Altered expression of a-type but not b-type synapsin isoform in the brain of patients at high risk for Alzheimer’s disease assessed by DNA microarray technique. Neurosci. Lett. 298, 191–194 (2001).

21. Emilsson, L., Saetre, P. & Jazin, E. Alzheimer’s disease: mRNA expression profiles of multiple patients show alterations of genes involved with calcium signaling. Neurobiol. Dis. 21, 618–625 (2006).

22. Xu, P.-T. et al. Differences in apolipoprotein E3/3 and E4/4 allele-specific gene expression in hippocampus in Alzheimer disease. Neurobiol. Dis. 21, 256–275 (2006).

23. Haroutunian, V., Katsel, P. & Schmeidler, J. Transcriptional vulnerability of brain regions in Alzheimer’s disease and dementia. Neurobiol. Aging 30, 561–573 (2009).

24. Weeraratna, A. T. et al. Alterations in immunological and neurological gene expression patterns in Alzheimer’s disease tissues. Exp. Cell Res. 313, 450–461 (2007).

25. Blalock, E. M. et al. Gene microarrays in hippocampal aging: statistical profiling identifies novel processes correlated with cognitive impairment. J. Neurosci. Off. J. Soc. Neurosci. 23, 3807–3819 (2003).

26. Parachikova, A. et al. Inflammatory changes parallel the early stages of Alzheimer disease. Neurobiol. Aging 28, 1821–1833 (2007).

27. Blalock, E. M., Buechel, H. M., Popovic, J., Geddes, J. W. & Landfield, P. W. Microarray analyses of laser-captured hippocampus reveal distinct gray and white matter signatures associated with incipient Alzheimer’s disease. J. Chem. Neuroanat. 42, 118–126 (2011).

28. Stark, R., Grzelak, M. & Hadfield, J. RNA sequencing: the teenage years. Nat. Rev. Genet. 20, 631–656 (2019).

29. Thorsson, V. et al. The Immune Landscape of Cancer. Immunity 48, 812–830.e14 (2018).

30. Northcott, P. A. et al. The whole-genome landscape of medulloblastoma subtypes. Nature 547, 311–317 (2017).

31. Gandal, M. J. et al. Transcriptome-wide isoform-level dysregulation in ASD, schizophrenia, and bipolar disorder. Science 362, eaat8127 (2018).

32. Marques-Coelho, D. et al. Differential transcript usage unravels gene expression alterations in Alzheimer’s disease human brains. Npj Aging Mech. Dis. 7, 2 (2021).

33. AD Knowledge Portal. https://adknowledgeportal.synapse.org/.

34. Allen, M. et al. Human whole genome genotype and transcriptome data for Alzheimer’s and other neurodegenerative diseases. Sci. Data 3, 160089 (2016).

35. Wang, M. et al. The Mount Sinai cohort of large-scale genomic, transcriptomic and proteomic data in Alzheimer’s disease. Sci. Data 5, 180185 (2018).

36. De Jager, P. L. et al. A multi-omic atlas of the human frontal cortex for aging and Alzheimer’s disease research. Sci. Data 5, 180142 (2018).

37. Willer, C. J., Li, Y. & Abecasis, G. R. METAL: fast and efficient meta-analysis of genomewide association scans. Bioinformatics 26, 2190 (2010).

38. Zhou, Y. et al. Metascape provides a biologist-oriented resource for the analysis of systems-level datasets. Nat. Commun. 10, 1523 (2019).

39. Kurtzer, G. M., Sochat, V. & Bauer, M. W. Singularity: Scientific containers for mobility of compute. PLoS ONE 12, e0177459 (2017).

40. Page, M. J. et al. The PRISMA 2020 statement: an updated guideline for reporting systematic reviews. BMJ 372, n71 (2021).

41. Raj, T. et al. Integrative transcriptome analyses of the aging brain implicate altered splicing in Alzheimer’s disease susceptibility. Nat. Genet. 50, 1584–1592 (2018).

42. Panitch, R. et al. Integrative brain transcriptome analysis links complement component 4 and HSPA2 to the APOE ε2 protective effect in Alzheimer disease. Mol. Psychiatry 26, 6054–6064 (2021).

43. Li, Q. S. & De Muynck, L. Differentially expressed genes in Alzheimer’s disease highlighting the roles of microglia genes including OLR1 and astrocyte gene CDK2AP1. Brain Behav. Immun. - Health 13, 100227 (2021).

44. Luo, D. et al. Integrative Transcriptomic Analyses of Hippocampal–Entorhinal System Subfields Identify Key Regulators in Alzheimer’s Disease. Adv. Sci. 10, 2300876 (2023).

45. Miller, J. A. et al. Neuropathological and transcriptomic characteristics of the aged brain. eLife 6, e31126 (2017).

46. Twine, N. A., Janitz, K., Wilkins, M. R. & Janitz, M. Whole Transcriptome Sequencing Reveals Gene Expression and Splicing Differences in Brain Regions Affected by Alzheimer’s Disease. PLoS ONE 6, e16266 (2011).

47. Mills, J. D. et al. RNA-Seq analysis of the parietal cortex in Alzheimer’s disease reveals alternatively spliced isoforms related to lipid metabolism. Neurosci. Lett. 536, 90–95 (2013).

48. Jia, Y. et al. Proteomic and Transcriptomic Analyses Reveal Pathological Changes in the Entorhinal Cortex Region that Correlate Well with Dysregulation of Ion Transport in Patients with Alzheimer’s Disease. Mol. Neurobiol. 58, 4007– 4027 (2021).

49. Shmookler Reis, R. J., et al. “Protein aggregates” contain RNA and DNA, entrapped by misfolded proteins but largely rescued by slowing translational elongation. Aging Cell 20, e13326 (2021).

50. Van Rooij, J. G. J. et al. Hippocampal transcriptome profiling combined with protein-protein interaction analysis elucidates Alzheimer’s disease pathways and genes. Neurobiol. Aging 74, 225–233 (2019).

51. Santana, D. A. et al. The Role of H3K9 Acetylation and Gene Expression in Different Brain Regions of Alzheimer’s Disease Patients. Epigenomics 14, 651–670 (2022).

52. Piras, I. S. et al. Association of AEBP1 and NRN1 RNA expression with Alzheimer’s disease and neurofibrillary tangle density in middle temporal gyrus. Brain Res. 1719, 217–224 (2019).

53. Magistri, M., Velmeshev, D., Makhmutova, M. & Faghihi, M. A. Transcriptomics Profiling of Alzheimer’s Disease Reveal Neurovascular Defects, Altered Amyloid-β Homeostasis, and Deregulated Expression of Long Noncoding RNAs. J. Alzheimers Dis. 48, 647–665 (2015).

54. Lee, M. Y. et al. Epigenome signatures landscaped by histone H3K9me3 are associated with the synaptic dysfunction in Alzheimer’s disease. Aging Cell 19, e13153 (2020).

55. King, D. et al. Synaptic resilience is associated with maintained cognition during ageing. Alzheimers Dement. 19, 2560–2574 (2023).

56. Humphries, C. E. et al. Integrated Whole Transcriptome and DNA Methylation Analysis Identifies Gene Networks Specific to Late-Onset Alzheimer’s Disease. J. Alzheimers Dis. 44, 977–987 (2015).

57. Guennewig, B. et al. Defining early changes in Alzheimer’s disease from RNA sequencing of brain regions differentially affected by pathology. Sci. Rep. 11, 4865 (2021).

58. Fisher, D. W., Tulloch, J., Yu, C.-E. & Tsuang, D. A Preliminary Comparison of the Methylome and Transcriptome from the Prefrontal Cortex Across Alzheimer’s Disease and Lewy Body Dementia. J. Alzheimers Dis. Rep. 7, 279–297 (2023).

59. Annese, A. et al. Whole transcriptome profiling of Late-Onset Alzheimer’s Disease patients provides insights into the molecular changes involved in the disease. Sci. Rep. 8, 4282 (2018).

60. Das, S. et al. Distinct transcriptomic responses to Aβ plaques, neurofibrillary tangles, and *APOE* in Alzheimer’s disease. Alzheimers Dement. 20, 74–90 (2024).

61. Felsky, D. et al. The Caribbean-Hispanic Alzheimer’s disease brain transcriptome reveals ancestry-specific disease mechanisms. Neurobiol. Dis. 176, 105938 (2023).

62. Cho, H. et al. Alterations of transcriptome signatures in head trauma-related neurodegenerative disorders. Sci. Rep. 10, 8811 (2020).

63. Caldwell, A. B. et al. Transcriptomic profiling of sporadic Alzheimer’s disease patients. Mol. Brain 15, 83 (2022).

64. Barbash, S. et al. Alzheimer’s brains show inter-related changes in RNA and lipid metabolism. Neurobiol. Dis. 106, 1–13 (2017).

65. Love, M. I., Soneson, C. & Patro, R. Swimming downstream: statistical analysis of differential transcript usage following Salmon quantification. F1000Research **7**, 952 (2018).

66. Aguzzoli Heberle, B., et al. Mapping medically relevant RNA isoform diversity in the aged human frontal cortex with deep long-read RNA-seq. Nat. Biotechnol. 1–12 (2024) doi:10.1038/s41587-024-02245-9.

67. Dong, X. et al. Benchmarking long-read RNA-sequencing analysis tools using in silico mixtures. Nat. Methods 20, 1810–1821 (2023).

68. Pardo-Palacios, F. J. et al. Systematic assessment of long-read RNA-seq methods for transcript identification and quantification. Nat. Methods 21, 1349–1363 (2024).

69. Chen, Y. et al. A systematic benchmark of Nanopore long read RNA sequencing for transcript level analysis in human cell lines. 2021.04.21.440736 Preprint at 10.1101/2021.04.21.440736 (2021).

70. Lu, G. et al. Protein kinase D 3 is localized in vesicular structures and interacts with vesicle-associated membrane protein 2. Cell. Signal. 19, 867–879 (2007).

71. Nelson, A. R., Sagare, A. P. & Zlokovic, B. V. Role of clusterin in the brain vascular clearance of amyloid-β. Proc. Natl. Acad. Sci. U. S. A. 114, 8681–8682 (2017).

72. Zhou, J. et al. BACE1 regulates expression of Clusterin in astrocytes for enhancing clearance of β-amyloid peptides. Mol. Neurodegener. 18, 31 (2023).

73. Pittaluga, A., Roggeri, A., Vallarino, G. & Olivero, G. Somatostatin, a Presynaptic Modulator of Glutamatergic Signal in the Central Nervous System. Int. J. Mol. Sci. 22, 5864 (2021).

74. Powell, A. J. et al. Recognition of structurally diverse substrates by type II 3-hydroxyacyl-CoA dehydrogenase (HADH II)/amyloid-beta binding alcohol dehydrogenase (ABAD). J. Mol. Biol. 303, 311–327 (2000).

75. Shi, G. et al. Functional alteration of PARL contributes to mitochondrial dysregulation in Parkinson’s disease. Hum. Mol. Genet. 20, 1966–1974 (2011).

76. Roberts, E. & Frankel, S. Glutamic acid decarboxylase in brain. J. Biol. Chem. 188, 789–795 (1951).

77. Roberts, E. & Frankel, S. gamma-Aminobutyric acid in brain: its formation from glutamic acid. J. Biol. Chem. 187, 55–63 (1950).

78. Han, Y. et al. STEAP3 Affects Ovarian Cancer Progression by Regulating Ferroptosis through the p53/SLC7A11 Pathway. Mediators Inflamm. 2024, 4048527 (2024).

79. Ye, C. L. et al. STEAP3 Affects Ferroptosis and Progression of Renal Cell Carcinoma Through the p53/xCT Pathway. Technol. Cancer Res. Treat. 21, 15330338221078728 (2022).

80. Bourgeois, B. & Madl, T. Regulation of cellular senescence via the FOXO4-p53 axis. FEBS Lett. 592, 2083–2097 (2018).

81. Orea-Soufi, A. et al. FOXO transcription factors as therapeutic targets in human diseases. Trends Pharmacol. Sci. 43, 1070–1084 (2022).

82. Dimova, E. Y., Samoylenko, A. & Kietzmann, T. FOXO4 induces human plasminogen activator inhibitor-1 gene expression via an indirect mechanism by modulating HIF-1alpha and CREB levels. Antioxid. Redox Signal. 13, 413–424 (2010).

83. Tang, T. T.-L. & Lasky, L. A. The forkhead transcription factor FOXO4 induces the down-regulation of hypoxia-inducible factor 1 alpha by a von Hippel-Lindau protein-independent mechanism. J. Biol. Chem. 278, 30125–30135 (2003).

84. Grubman, A. et al. Transcriptional signature in microglia associated with Aβ plaque phagocytosis. Nat. Commun. 12, 3015 (2021).

85. Baik, S. H. et al. A Breakdown in Metabolic Reprogramming Causes Microglia Dysfunction in Alzheimer’s Disease. Cell Metab. 30, 493–507.e6 (2019).

86. Chou, V. et al. INPP5D regulates inflammasome activation in human microglia. Nat. Commun. 14, 7552 (2023).

87. Kijima, K. et al. Periaxin mutation causes early-onset but slow-progressive Charcot-Marie-Tooth disease. J. Hum. Genet. 49, 376–379 (2004).

88. Brenner, D. et al. Hot-spot KIF5A mutations cause familial ALS. Brain 141, 688–697 (2018).

89. Soustelle, L. et al. ALS-Associated KIF5A Mutation Causes Locomotor Deficits Associated with Cytoplasmic Inclusions, Alterations of Neuromuscular Junctions, and Motor Neuron Loss. J. Neurosci. 43, 8058–8072 (2023).

90. Mittelstaedt, T., Alvaréz-Baron, E. & Schoch, S. RIM proteins and their role in synapse function. 391, 599–606 (2010).

91. Bonham, L. W. et al. CXCR4 involvement in neurodegenerative diseases. Transl. Psychiatry 8, 1–10 (2018).

92. Sánchez-Juan, P. et al. Serum GFAP levels correlate with astrocyte reactivity, post-mortem brain atrophy and neurofibrillary tangles. Brain J. Neurol. 147, 1667–1679 (2024).

93. Brenner, M. Role of GFAP in CNS injuries. Neurosci. Lett. 565, 7–13 (2014).

94. Kim, K. Y., Shin, K. Y. & Chang, K.-A. GFAP as a Potential Biomarker for Alzheimer’s Disease: A Systematic Review and Meta-Analysis. Cells 12, 1309 (2023).

95. Bellaver, B. et al. Astrocyte Biomarkers in Alzheimer Disease. Neurology 96, e2944–e2955 (2021).

96. Ferrari-Souza, J. P. et al. Astrocyte biomarker signatures of amyloid-β and tau pathologies in Alzheimer’s disease. Mol. Psychiatry 27, 4781–4789 (2022).

97. Holper, S. et al. Blood Astrocyte Biomarkers in Alzheimer Disease. Neurology 103, e209537 (2024).

98. Benedet, A. L. et al. Differences Between Plasma and Cerebrospinal Fluid Glial Fibrillary Acidic Protein Levels Across the Alzheimer Disease Continuum. JAMA Neurol. 78, 1–13 (2021).

99. Alqarni, S. & Alsebai, M. Could VGF and/or its derived peptide act as biomarkers for the diagnosis of neurodegenerative diseases: A systematic review. Front. Endocrinol. 13, 1032192 (2022).

100. Quinn, J. P., Kandigian, S. E., Trombetta, B. A., Arnold, S. E. & Carlyle, B. C. VGF as a biomarker and therapeutic target in neurodegenerative and psychiatric diseases. Brain Commun. 3, fcab261 (2021).

101. Bennett, G. W., Ballard, T. M., Watson, C. D. & Fone, K. C. Effect of neuropeptides on cognitive function. Exp. Gerontol. 32, 451–469 (1997).

102. Belloy, M. E. et al. APOE Genotype and Alzheimer Disease Risk Across Age, Sex, and Population Ancestry. JAMA Neurol. 80, 1284–1294 (2023).

103. Keren-Shaul, H. et al. A Unique Microglia Type Associated with Restricting Development of Alzheimer’s Disease. Cell 169, 1276–1290.e17 (2017).

104. Mathys, H. et al. Single-cell atlas reveals correlates of high cognitive function, dementia, and resilience to Alzheimer’s disease pathology. Cell 186, 4365–4385.e27 (2023).

105. Mathys, H. et al. Single-cell multiregion dissection of Alzheimer’s disease. Nature 1–11 (2024) doi:10.1038/s41586-024-07606-7.

106. Sadick, J. S. et al. Astrocytes and oligodendrocytes undergo subtype-specific transcriptional changes in Alzheimer’s disease. Neuron 110, 1788–1805.e10 (2022).

107. Mathys, H. et al. Single-cell transcriptomic analysis of Alzheimer’s disease. Nature 570, 332–337 (2019).

108. Zhou, Y. et al. Human and mouse single-nucleus transcriptomics reveal TREM2-dependent and TREM2- independent cellular responses in Alzheimer’s disease. Nat. Med. 26, 131–142 (2020).

109. Grubman, A. et al. A single-cell atlas of entorhinal cortex from individuals with Alzheimer’s disease reveals cell-type-specific gene expression regulation. Nat. Neurosci. 22, 2087–2097 (2019).

110. Chen, S. et al. Spatially resolved transcriptomics reveals genes associated with the vulnerability of middle temporal gyrus in Alzheimer’s disease. Acta Neuropathol. Commun. 10, 188 (2022).

111. Zhao, B. S., Roundtree, I. A. & He, C. Post-transcriptional gene regulation by mRNA modifications. Nat. Rev. Mol. Cell Biol. 18, 31–42 (2017).

112. Furlan, M. et al. Computational methods for RNA modification detection from nanopore direct RNA sequencing data. RNA Biol. 18, 31–40 (2021).

113. van Dijk, E. L. et al. Genomics in the long-read sequencing era. Trends Genet. TIG 39, 649–671 (2023).

114. Zhao, X., Zhang, Y., Hang, D., Meng, J. & Wei, Z. Detecting RNA modification using direct RNA sequencing: A systematic review. Comput. Struct. Biotechnol. J. 20, 5740–5749 (2022).

115. Cerneckis, J., Ming, G.-L., Song, H., He, C. & Shi, Y. The rise of epitranscriptomics: recent developments and future directions. Trends Pharmacol. Sci. 45, 24–38 (2024).

116. Sağlam, B. & Akgül, B. An Overview of Current Detection Methods for RNA Methylation. Int. J. Mol. Sci. 25, 3098 (2024).

