## Supplementary Figures for "Systematic review and meta-analysis of bulk RNAseq studies in human Alzheimer’s disease brain tissue"

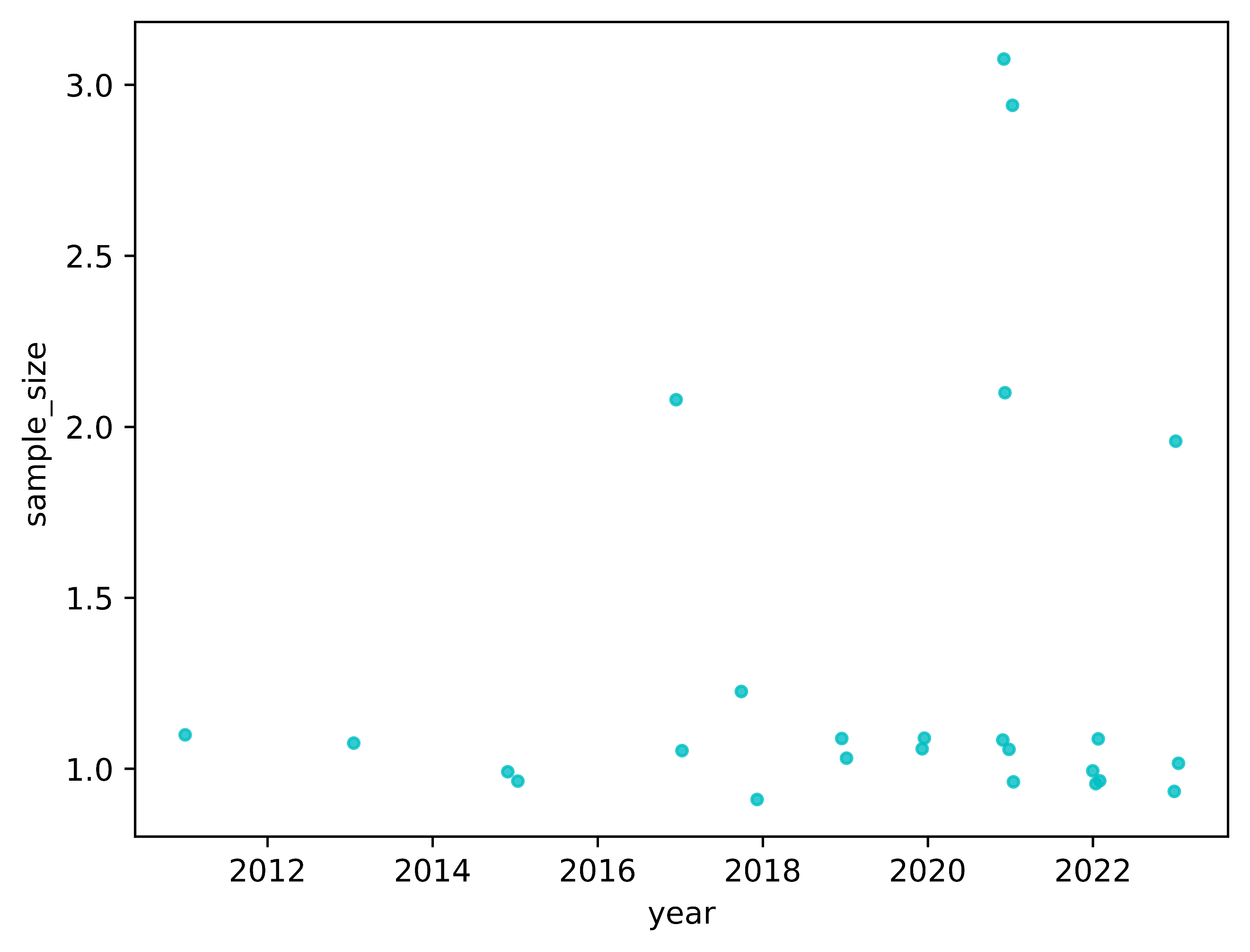

**Supplementary Figure 1:** The scatterplot illustrates the quality assessment score for sample size on the Y-axis and the year of study publication on the X-axis. Spearman coefficient = 0.09 and p-value = 0.68. The correlation did not reach statistical significance after Bonferroni correction, with a threshold for significance set at Bonferroni corrected p-value < 0.1 (unadjusted p-value threshold < 0.009). Jitter was added to the data points to reduce overlap and enhance visual clarity, without affecting the Spearman coefficient and p-value calculations.

­
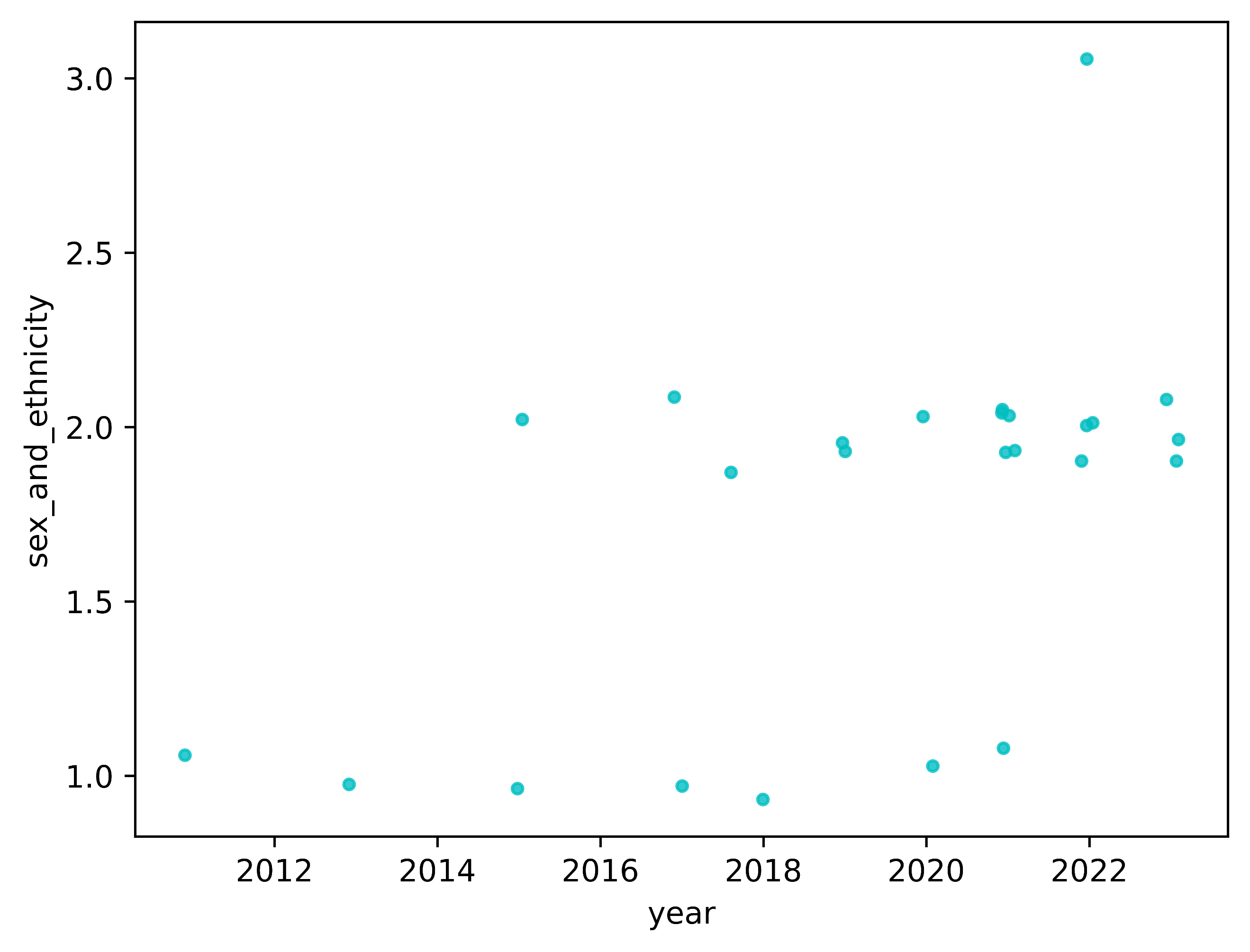

**Supplementary Figure 2:** The scatterplot illustrates the quality assessment score for sex and ethnicity on the Y-axis and the year of study publication on the X-axis. Spearman coefficient = 0.61 and p-value = 0.0012. The correlation reached statistical significance after Bonferroni correction, with a threshold for significance set at Bonferroni corrected p-value < 0.1 (unadjusted p-value threshold < 0.009). Jitter was added to the data points to reduce overlap and enhance visual clarity, without affecting the Spearman coefficient and p-value calculations.

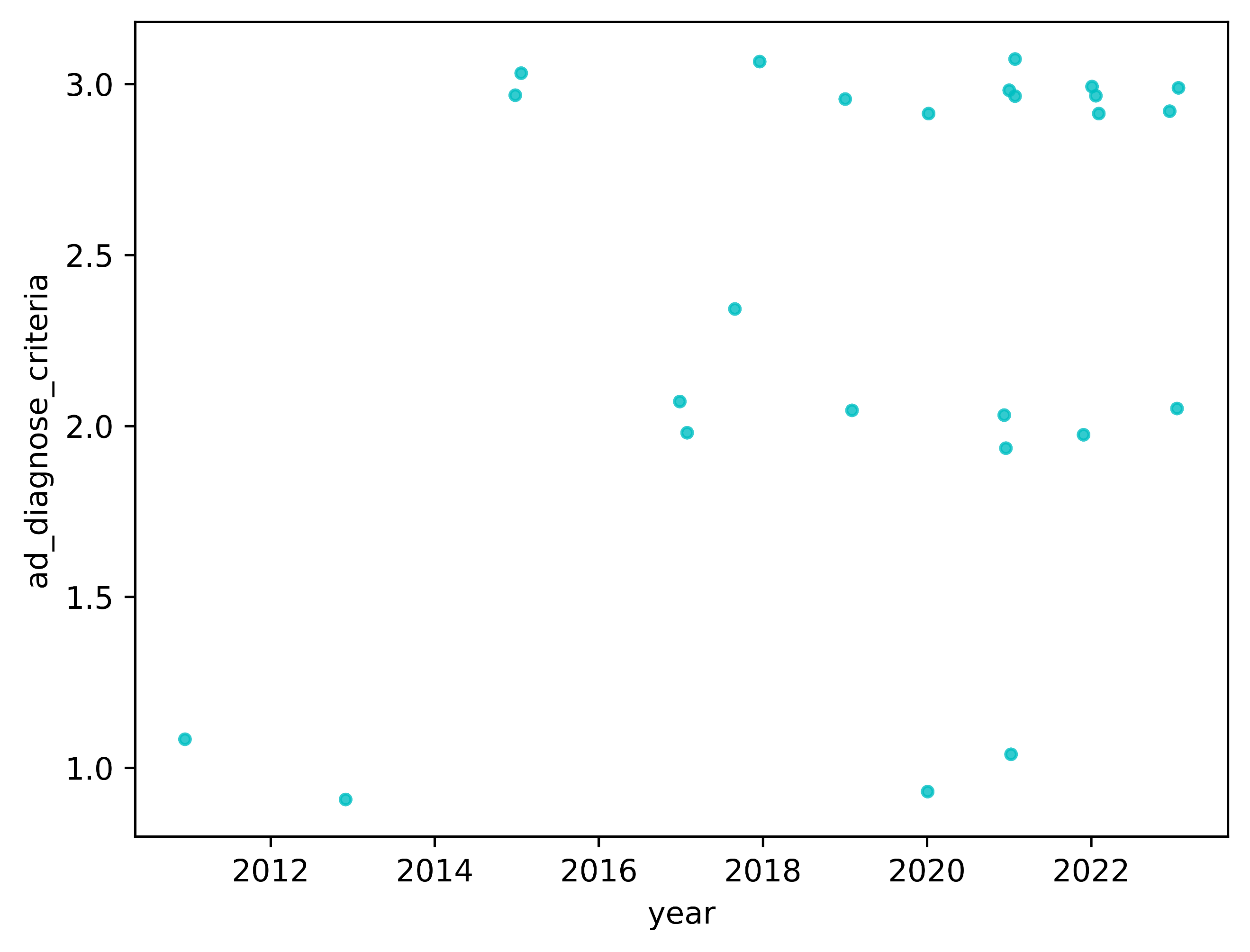

**Supplementary Figure 3:** The scatterplot illustrates the quality assessment score for AD diagnose criteria on the Y-axis and the year of study publication on the X-axis. Spearman coefficient = 0.3 and p-value = 0.15. The correlation did not statistical significance after Bonferroni correction, with a threshold for significance set at Bonferroni corrected p-value < 0.1 (unadjusted p-value threshold < 0.009). Jitter was added to the data points to reduce overlap and enhance visual clarity, without affecting the Spearman coefficient and p-value calculations.

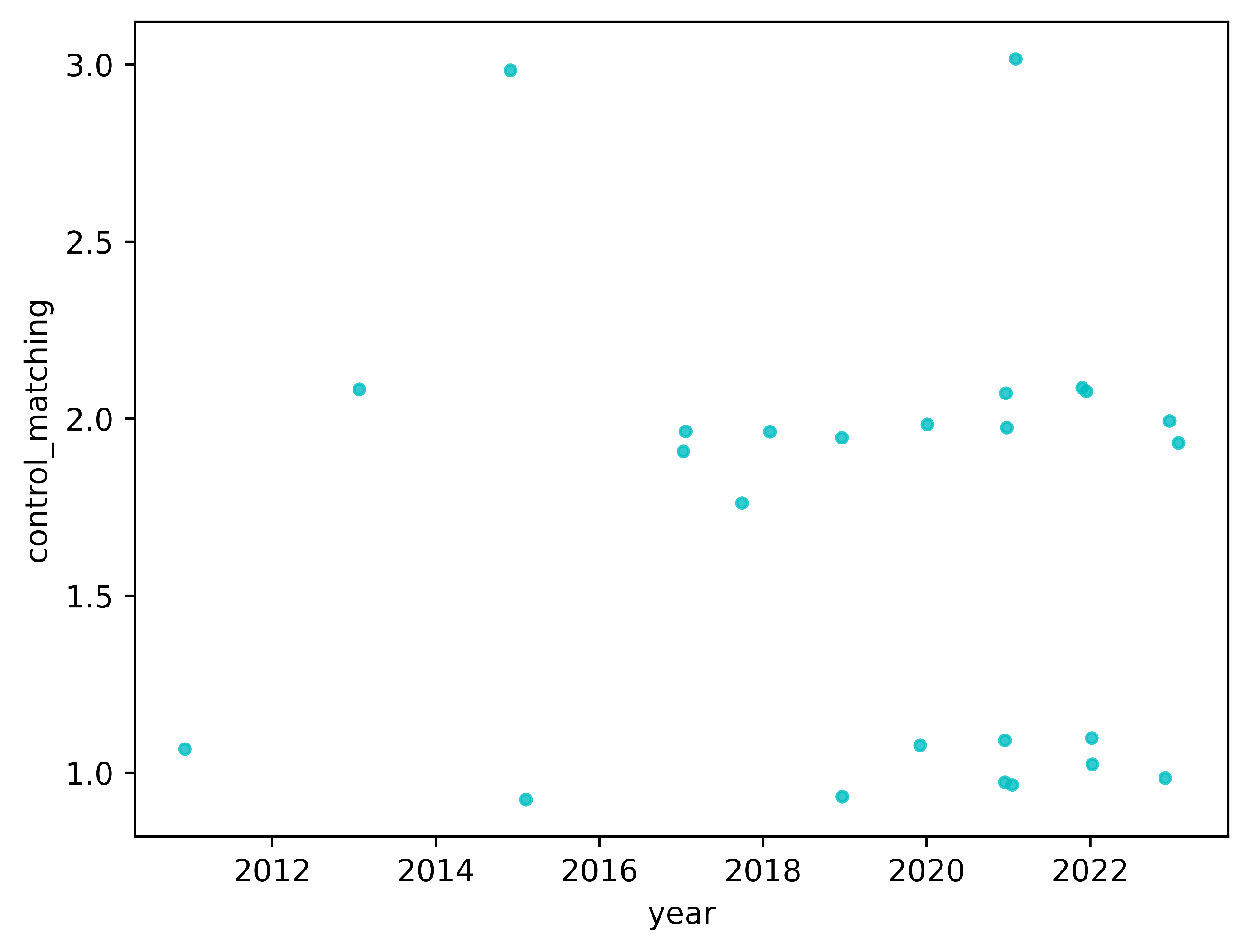

**Supplementary Figure 4:** The scatterplot illustrates the quality assessment score for control matching on the Y-axis and the year of study publication on the X-axis. Spearman coefficient = -0.08 and p-value = 0.70. The correlation did not reach statistical significance after Bonferroni correction, with a threshold for significance set at Bonferroni corrected p-value < 0.1 (unadjusted p-value threshold < 0.009). Jitter was added to the data points to reduce overlap and enhance visual clarity, without affecting the Spearman coefficient and p-value calculations.

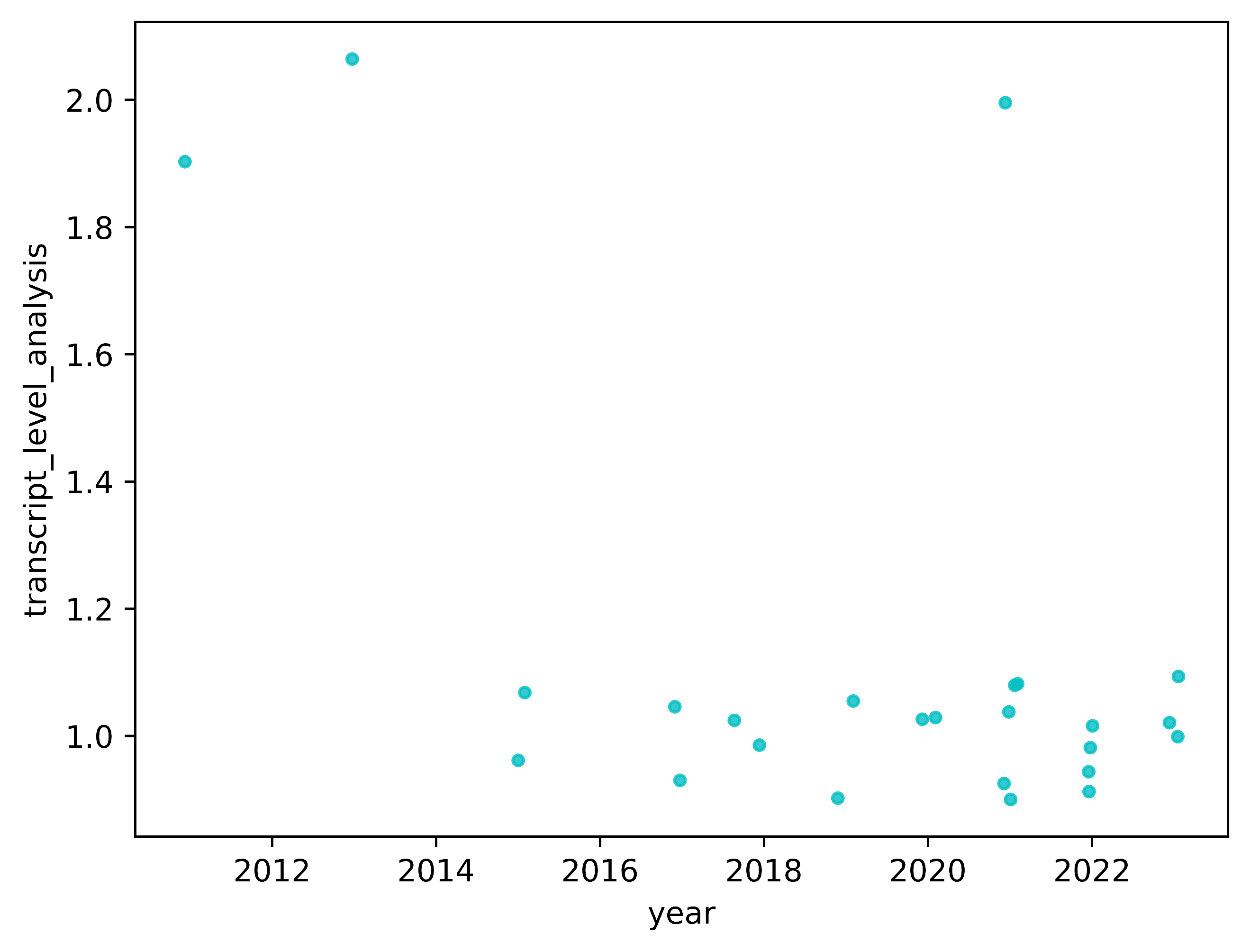

**Supplementary Figure 5:** The scatterplot illustrates the quality assessment score for transcript-level analysis on the Y-axis and the year of study publication on the X-axis. Spearman coefficient = -0.41 and p-value = 0.16. The correlation did not reach statistical significance after Bonferroni correction, with a threshold for significance set at Bonferroni corrected p-value < 0.1 (unadjusted p-value threshold < 0.009). Jitter was added to the data points to reduce overlap and enhance visual clarity, without affecting the Spearman coefficient and p-value calculations.

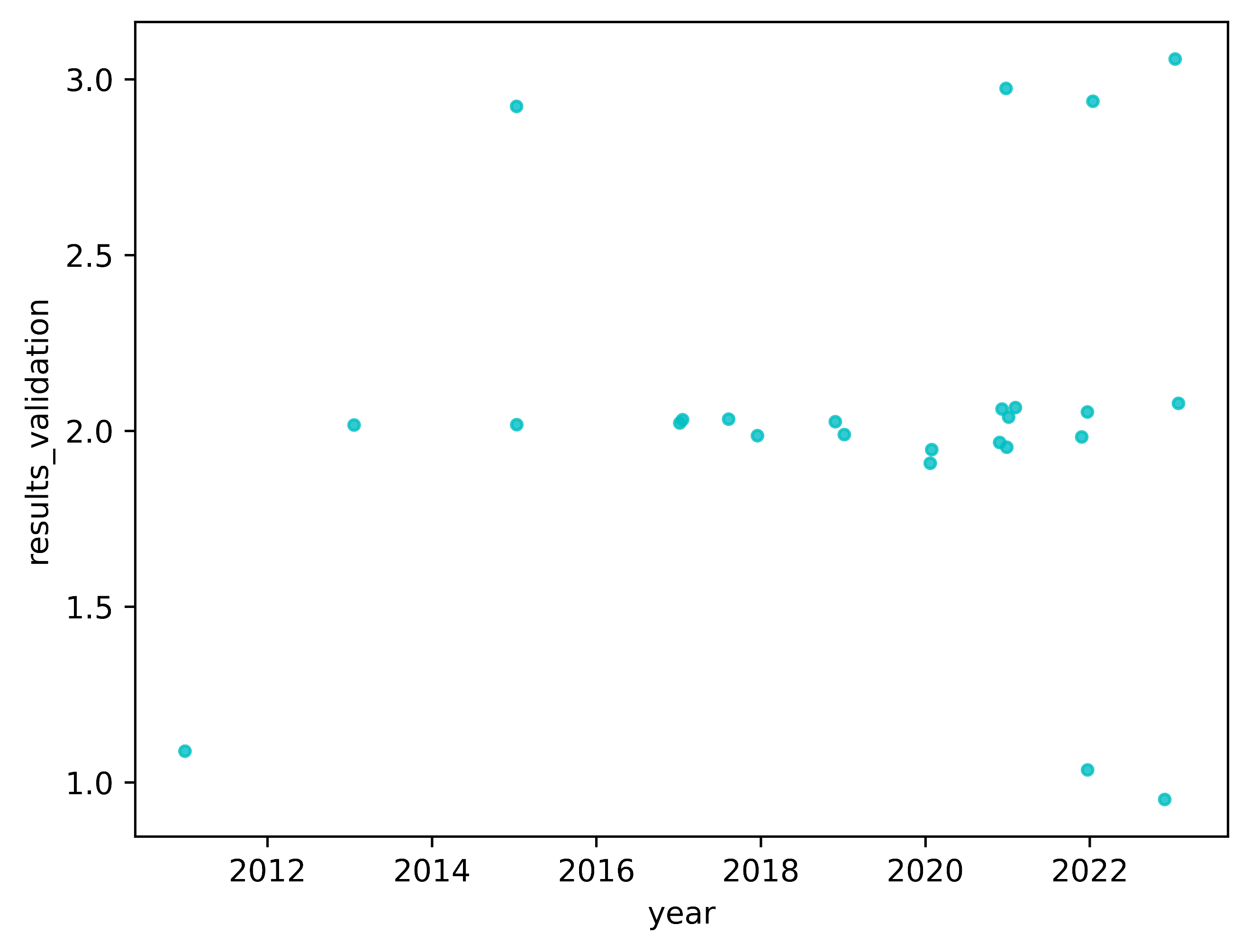

**Supplementary Figure 6:** The scatterplot illustrates the quality assessment score for results validation on the Y-axis and the year of study publication on the X-axis. Spearman coefficient = 0.06 and p-value = 0.79. The correlation did not reach statistical significance after Bonferroni correction, with a threshold for significance set at Bonferroni corrected p-value < 0.1 (unadjusted p-value threshold < 0.009). Jitter was added to the data points to reduce overlap and enhance visual clarity, without affecting the Spearman coefficient and p-value calculations.

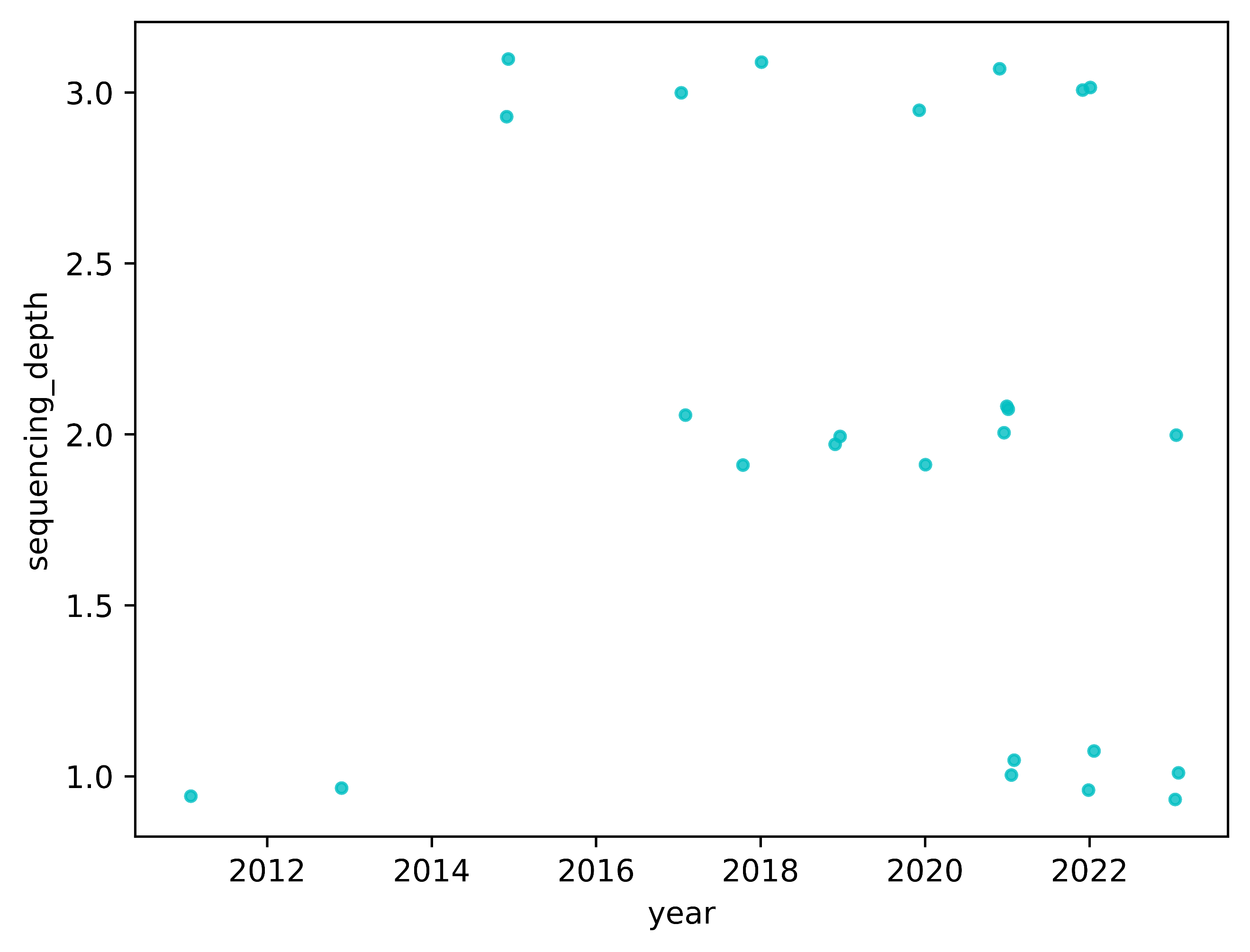

**Supplementary Figure 7:** The scatterplot illustrates the quality assessment score for sequencing depth on the Y-axis and the year of study publication on the X-axis. Spearman coefficient = -0.24 and p-value = 0.24. The correlation did not reach statistical significance after Bonferroni correction, with a threshold for significance set at Bonferroni corrected p-value < 0.1 (unadjusted p-value threshold < 0.009). Jitter was added to the data points to reduce overlap and enhance visual clarity, without affecting the Spearman coefficient and p-value calculations.

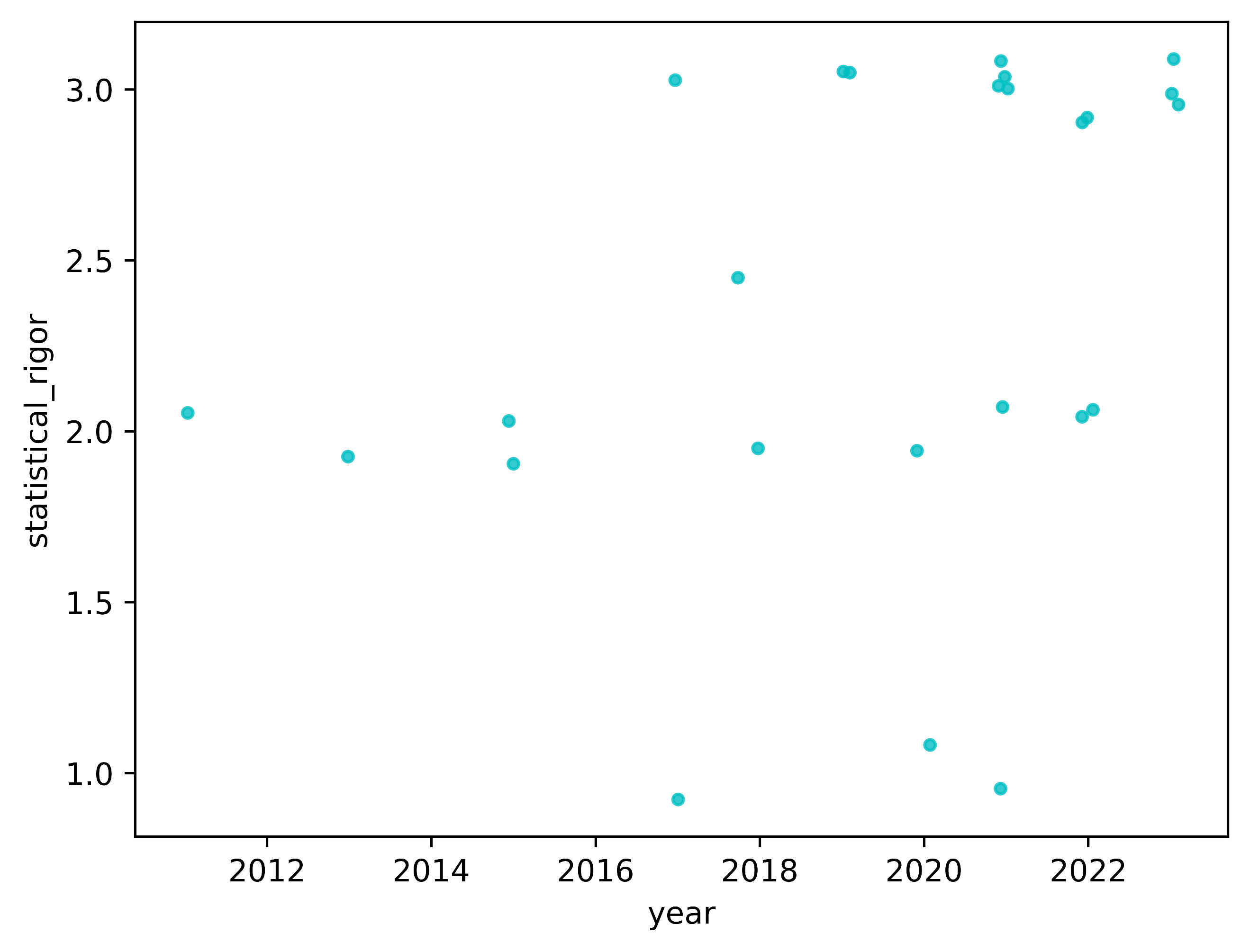

**Supplementary Figure 8:** The scatterplot illustrates the quality assessment score for statistical rigor on the Y-axis and the year of study publication on the X-axis. Spearman coefficient = 0.43 and p-value = 0.031. The correlation did not reach statistical significance after Bonferroni correction, with a threshold for significance set at Bonferroni corrected p-value < 0.1 (unadjusted p-value threshold < 0.009). Jitter was added to the data points to reduce overlap and enhance visual clarity, without affecting the Spearman coefficient and p-value calculations.

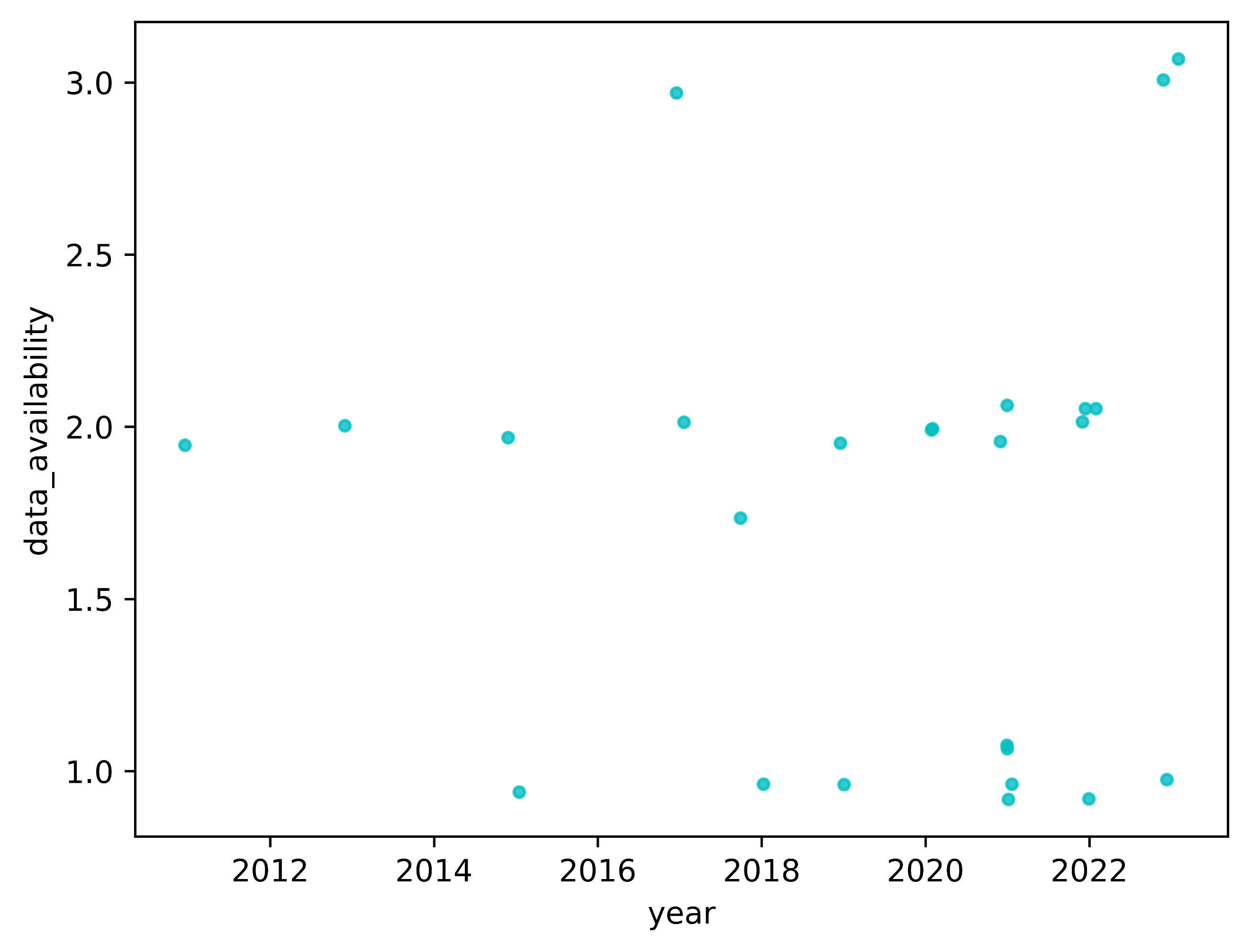

**Supplementary Figure 9:** The scatterplot illustrates the quality assessment score for data availability on the Y-axis and the year of study publication on the X-axis. Spearman coefficient = 0.02 and p-value = 0.91. The correlation did not reach statistical significance after Bonferroni correction, with a threshold for significance set at Bonferroni corrected p-value < 0.1 (unadjusted p-value threshold < 0.009). Jitter was added to the data points to reduce overlap and enhance visual clarity, without affecting the Spearman coefficient and p-value calculations.

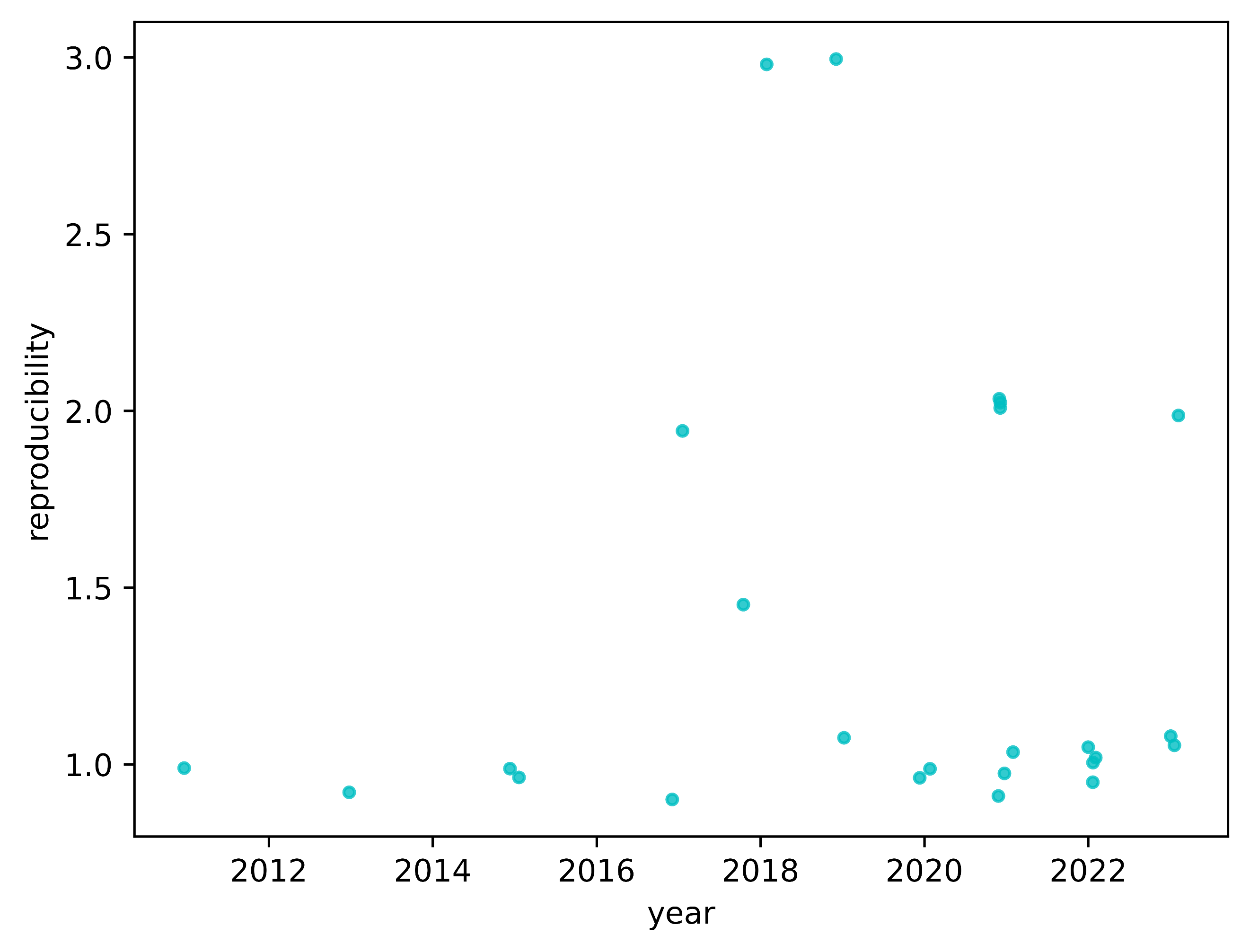

**Supplementary Figure 10:** The scatterplot illustrates the quality assessment score for reproducibility on the Y-axis and the year of study publication on the X-axis. Spearman coefficient = -0.05 and p-value = 0.81. The correlation did not reach statistical significance after Bonferroni correction, with a threshold for significance set at Bonferroni corrected p-value < 0.1 (unadjusted p-value threshold < 0.009). Jitter was added to the data points to reduce overlap and enhance visual clarity, without affecting the Spearman coefficient and p-value calculations.

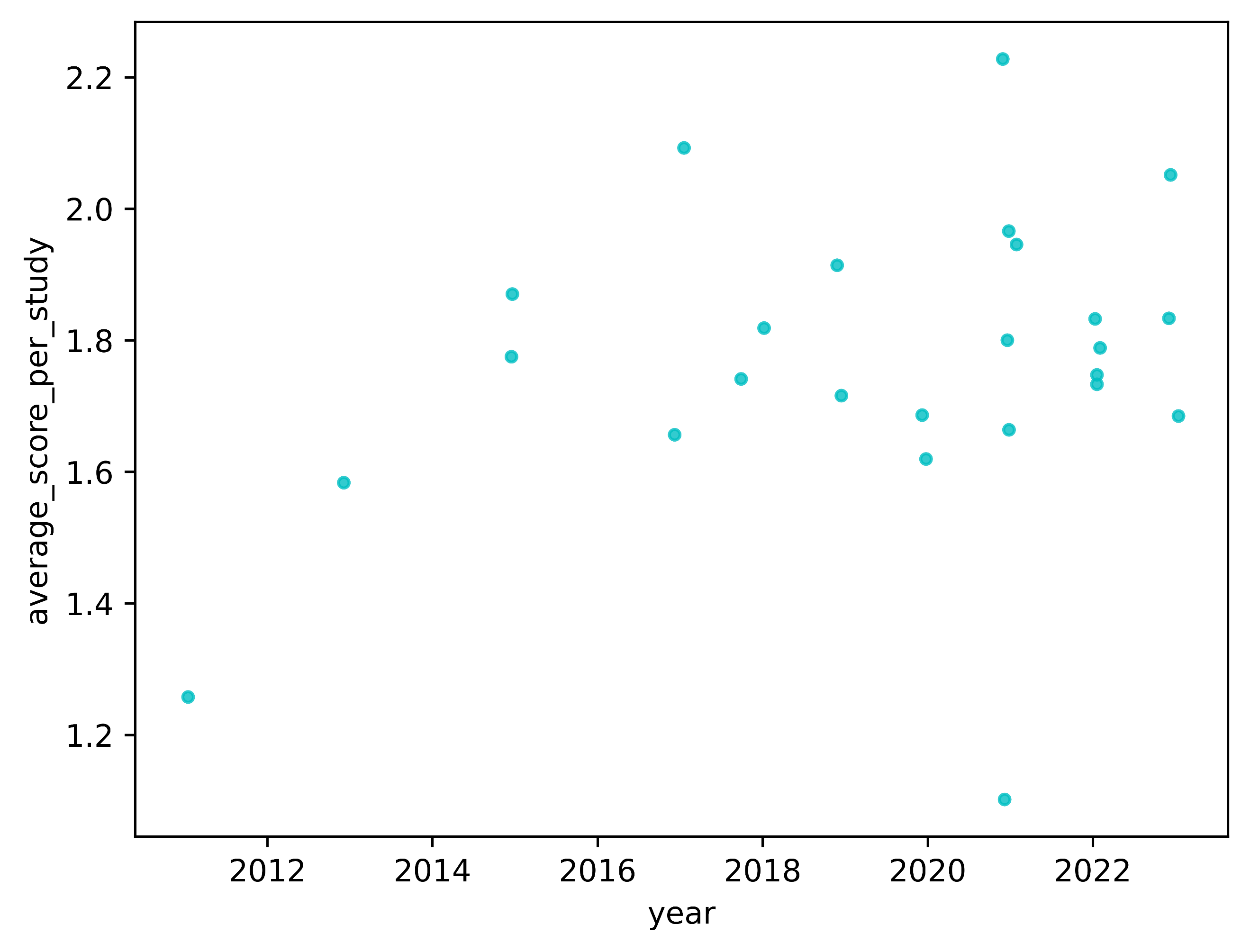

**Supplementary Figure 11:** The scatterplot illustrates the quality assessment score for average score per study on the Y-axis and the year of sudy publication on the X-axis. Spearman coefficient = 0.22 and p-value = 0.29. The correlation did not reach statistical significance after Bonferroni correction, with a threshold for significance set at Bonferroni corrected p-value < 0.1 (unadjusted p-value threshold < 0.009). Jitter was added to the data points to reduce overlap and enhance visual clarity, without affecting the Spearman coefficient and p-value calculations.

**Supplementary Table Descriptions:**

**Supplementary Table 1:** Definition of terms used for data extraction.

**Supplementary Table 2:** Data extraction table for studies included in the systematic review.

**Supplementary Table 3:** Software versions used for data analysis and visualizations in this article.

**Supplementary Table 4:** Correlations between year of publication and quality assessment category scores.

**Supplementary Table 5:** Full meta-analysis differential expression results for temporal lobe.

**Supplementary Table 6:** Genes upregulated in Alzheimer's

disease patients for the temporal lobe differential gene expression meta-analysis.

**Supplementary Table 7:** Genes downregulated in Alzheimer's

disease subjects for the temporal lobe differential gene expression meta-analysis.

**Supplementary Table 8:** Overlap between differentially expressed genes found in our temporal lobe meta-analysis and other studies temporal lobe studies included in our systematic review. "number_of_overlapping_studies" equal to 0 means the differentially expressed gene was unique to our meta-analysis.

**Supplementary Table 9:** Summary statistics for heterogeneity analysis (I-squared) for temporal lobe and frontal lobe differential gene expression meta-analysis.

**Supplementary Table 10:** Full meta-analysis differential expression results for frontal lobe.

**Supplementary Table 11:** Genes upregulated in Alzheimer's

disease patients for the frontal lobe differential gene expression meta-analysis.

**Supplementary Table 12:** Genes downregulated in Alzheimer's

disease subjects for the frontal lobe differential gene expression meta-analysis.

**Supplementary Table 13:** Overlap between differentially expressed genes found in our frontal lobe meta-analysis and other studies frontal lobe studies included in our systematic review. "number_of_overlapping_studies" equal to 0 means the differentially expressed gene was unique to our meta-analysis.

**Supplementary Table 14:** Overlapping genes between temporal and frontal lobe meta-analysis.
