## Supplementary File 2 - Prisma Abstract Checklist for "Systematic review and meta-analysis of bulk RNAseq studies in human Alzheimer’s disease brain tissue"

| **Section and Topic** | **Item #** | **Checklist item** | **Reported (Yes/No)** |
| --- | --- | --- | --- |
| **TITLE** | | |  |
| Title | 1 | Identify the report as a systematic review. | Yes |
| **BACKGROUND** | | |  |
| Objectives | 2 | Provide an explicit statement of the main objective(s) or question(s) the review addresses. | Yes |
| **METHODS** | | |  |
| Eligibility criteria | 3 | Specify the inclusion and exclusion criteria for the review. | Yes |
| Information sources | 4 | Specify the information sources (e.g. databases, registers) used to identify studies and the date when each was last searched. | Yes |
| Risk of bias | 5 | Specify the methods used to assess risk of bias in the included studies. | Yes |
| Synthesis of results | 6 | Specify the methods used to present and synthesise results. | Yes |
| **RESULTS** | | |  |
| Included studies | 7 | Give the total number of included studies and participants and summarise relevant characteristics of studies. | Yes |
| Synthesis of results | 8 | Present results for main outcomes, preferably indicating the number of included studies and participants for each. If meta-analysis was done, report the summary estimate and confidence/credible interval. If comparing groups, indicate the direction of the effect (i.e. which group is favoured). | Yes |
| **DISCUSSION** | | |  |
| Limitations of evidence | 9 | Provide a brief summary of the limitations of the evidence included in the review (e.g. study risk of bias, inconsistency and imprecision). | Yes |
| Interpretation | 10 | Provide a general interpretation of the results and important implications. | Yes |
| **OTHER** | | |  |
| Funding | 11 | Specify the primary source of funding for the review. | Yes |
| Registration | 12 | Provide the register name and registration number. | Yes |

*From:*  Page MJ, McKenzie JE, Bossuyt PM, Boutron I, Hoffmann TC, Mulrow CD, et al. The PRISMA 2020 statement: an updated guideline for reporting systematic reviews. BMJ 2021;372:n71. doi: 10.1136/bmj.n71
